# A hierarchical clock-mixture model for Bayesian phylogenetic dating

**DOI:** 10.64898/2026.07.15.738815

**Authors:** Yuan Xu, Jordan Douglas, Remco R. Bouckaert, Alexei J. Drummond

**Affiliations:** Centre for Computational Evolution, University of Auckland, Auckland, New Zealand; School of Biological Sciences, University of Auckland, Auckland, New Zealand; Research School of Biology, Australian National University, ACT, Australia; Department of Physics, University of Auckland, Auckland, New Zealand; School of Computer Science, University of Auckland, Auckland, New Zealand

## Abstract

Conditioning the inference of a Bayesian phylogenetic time tree on a single molecular clock model treats clock choice as fixed, even when support among plausible clock families is uncertain. When that assumption is wrong, estimated timescales and their uncertainty can be distorted. Existing practice usually addresses this by fitting strict, uncorrelated lognormal relaxed (UCLN), and autocorrelated clocks separately and comparing their marginal likelihoods, but this requires multiple computationally expensive model selection analyses. Here we introduce a hierarchical clock-mixture framework, implemented as the open-source `RelaxClockAveraging` package for BEAST 2, that separates two questions that are often conflated in clock comparison: whether branch-specific rate variation is needed at all, and, if it is, whether that variation is better described as uncorrelated or autocorrelated. The method averages analytically between strict and relaxed clocks at the top level and then compares UCLN and autocorrelated models within the relaxed class on a shared branch-rate vector, returning posterior probabilities for all three clock families together with model-averaged summaries for parameters shared across them. In stratified simulations, the generating clock family was retained in the 95% posterior model set in all replicates, while model-averaged estimates of the overall substitution rate, root age, and tree length remained accurate. On a DENV-4 benchmark, the mixture reproduced the higher-effort nested-sampling ranking of clock families while avoiding the extreme run-to-run variability of independent marginal-likelihood estimates. On empirical benchmarks, the method recovered strong support for the autocorrelated clock on the classical 31-taxon chloroplast *rbcL* data set. On an RSV-A G-gene data set it concentrated virtually all posterior support on UCLN while preserving the established RSV-A timescale. On a 245-taxon structured-coalescent H3N2 dataset it assigned most posterior mass to the relaxed-clock class, with support within that class concentrated on the autocorrelated family, and the inferred timescale remained consistent with the published estimate. These results show that clock-model support and downstream timescale sensitivity are related but not identical. Single-clock dating analyses can still be adequate, but fixing a clock model should be justified by posterior support rather than treated as a default assumption.

## 1 Introduction

Absolute divergence times provide the temporal framework for asking when lineages diversified, when key traits evolved, and how biological radiations relate to geological and climatic change. In Bayesian molecular dating, these questions are addressed by combining sequence data with fossil or other temporal calibrations under explicit models of sequence evolution, branching times, and rate variation (Yang and Rannala, 2006; Drummond et al., 2006; Dos Reis et al., 2016; Reis et al., 2018). A central difficulty is that molecular data identify branch lengths in substitutions per site, whereas rates and times cannot be separated without external temporal information—such as fossil calibrations or, for measurably evolving populations, the sampling times of the sequences themselves—and assumptions about how rates vary across the tree (Drummond et al., 2002; Rannala, 2016). The clock model is therefore not a minor technical detail of a dating analysis. It determines how molecular change is partitioned into rate and time, and different clock assumptions can lead to different divergence-time estimates and different assessments of uncertainty (Duchêne et al., 2014; Ho and Duchêne, 2014; Dos Reis et al., 2016). This issue is not confined to fossil- or tip-dated analyses: even when only relative times, branch lengths, or downstream summaries such as ancestral-state reconstructions are of interest, the clock model can affect how sequence change is distributed across the tree (Cusimano and Renner, 2014; Duchêne et al., 2014; Ho and Duchêne, 2014).

The original molecular clock assumed that sequences accumulate substitutions at an approximately constant rate through time, and a strict clock can still be adequate when departures from clock-like evolution are limited (Zuckerkandl, 1962; Zuckerkandl and Pauling, 1965; Bromham and Penny, 2003; Ho and Duchêne, 2014). Many empirical data sets, however, show substantial among-lineage rate variation, reflecting differences in life history, mutation process, generation time, population size, and other biological factors (Bromham and Penny, 2003; Ho et al., 2015). Relaxed-clock models were introduced to accommodate this heterogeneity, but they do not all describe rate variation in the same way. Under the uncorrelated lognormal relaxed clock (UCLN), branch-specific rates are drawn independently from a common lognormal distribution governed by a log-scale dispersion parameter; as that dispersion approaches zero, the model approaches a strict clock (Drummond et al., 2006); under an autocorrelated clock, rates change along the tree so that descendant branches tend to resemble their ancestors (Thorne et al., 1998; Kishino et al., 2001). In this study, we focus on the three clock models that commonly define this choice in Bayesian phylogenetic dating—the strict clock, UCLN, and the autocorrelated clock—and treat clock choice as an inferential target rather than a default background setting (Lepage et al., 2007; Ho and Duchêne, 2014).

A common way to deal with this uncertainty is to fit competing clock models separately and compare their marginal likelihoods. In Bayesian terms this is a coherent strategy, and path-sampling, stepping-stone sampling, and nested sampling have all been used for phylogenetic model comparison (Lartillot and Philippe, 2006; Xie et al., 2011; Baele et al., 2012; Fan et al., 2011; Baele et al., 2016; Russel et al., 2019). In practice, however, these analyses are computationally demanding, can be sensitive to tuning, and become cumbersome once several clock models must each be fitted and compared separately. These difficulties have prompted a range of single-analysis alternatives, including model averaging across uncorrelated relaxed clocks (Li and Drummond, 2012; Douglas et al., 2025), random local clocks that infer a variable number of rate-change points across the tree within a single analysis (Drummond and Suchard, 2010), local relaxed clocks that fit a separate relaxed clock within each of a set of predefined tree regions delimited by fixed change points (Fourment and Darling, 2018), reversible-jump approaches for tip-dating that still do not include an autocorrelated clock (Zhang, 2022), analytical mixtures that allow competing prior models to contribute within a single Markov chain Monte Carlo (MCMC) analysis (Darlim and Höhna, 2024), and mixed relaxed-clock models that combine uncorrelated and autocorrelated behaviour within one generative process (Lartillot et al., 2016). These developments address parts of the broader problem, but none provides a single-analysis framework that jointly compares the strict clock, the UCLN clock, and an auto-correlated clock within one coherent Bayesian phylogenetic dating analysis. The analytical-mixture approach cannot include a strict clock because equal branch rates have zero probability under standard relaxed-clock priors (Darlim and Höhna, 2024), whereas mixed relaxed-clock models address a different question entirely: rather than choosing among discrete clock models, they define a new model that blends the two forms of relaxed-clock behaviour (Lartillot et al., 2016).

The unresolved problem is also conceptual. The three candidate clock models are not related to one another in the same way. A strict clock is the limiting case in which all branches are constrained to share a single rate, whereas both UCLN and autocorrelated clocks allow rates to vary among branches and differ mainly in how that variation is described across the tree (Drummond et al., 2006; Thorne et al., 1998; Kishino et al., 2001). Moving from a strict clock to a relaxed clock therefore introduces branch-specific rates that are absent under the strict clock, whereas moving from UCLN to an autocorrelated clock changes the assumptions about how those rates vary across the phylogeny. This makes a hierarchical treatment of clock uncertainty more natural than a flat comparison among three standalone models. Here we implement that idea in BEAST 2 (Bouckaert et al., 2019, 2014) by averaging between strict and relaxed clocks at the top level and, within the relaxed class, allowing support to be shared between UCLN and an autocorrelated clock defined on the same set of branch-specific rates. The result is a single analysis that returns posterior probabilities for all three clock models together with clock-averaged divergence-time estimates.

We evaluate the method in four steps. First, stratified simulations test whether posterior probabilities for the three clock models are well calibrated and whether carrying clock uncertainty through the analysis still allows accurate estimation of shared quantities such as the overall rate and root age. Second, we use a serially sampled dengue virus serotype 4 (DENV-4) data set as a practical benchmark against standalone nested sampling (Russel et al., 2019). Third, we reanalyse the classical data sets associated with the development of autocorrelated clocks to ask whether the method recovers the strong signal expected in that setting (Thorne et al., 1998). Finally, we analyse RSV-A and H3N2 data sets as contrasting modern applications, asking both whether the method recovers decisive support when one relaxed clock is clearly preferred and whether residual uncertainty within the relaxed class has a meaningful effect on the inferred timescale. These analyses let us ask not only which clock model is favoured, but also when it is more informative to carry clock uncertainty forward into the dating results.

## 2 Materials and Methods

### 2.1 Overview of the hierarchical clock model

The key design choice is to separate two questions that are often conflated in clock-model comparison. The first is whether substitution rates should be constrained to a single value across the tree or allowed to vary among branches. The second question arises only once branch-specific rate variation is allowed: should those rates be treated as independent draws, as in UCLN, or as correlated along the phylogeny, as in an autocorrelated clock? These two comparisons have different statistical structures. A strict clock asks whether branch-specific rate variation is needed at all, whereas the comparison between UCLN and an autocorrelated clock asks how that variation should be described once it is allowed.

This distinction determines the form of the hierarchical clock model. At the top level, the analysis averages between a strict-clock likelihood and a relaxed-clock likelihood. Within the relaxed part of the model, support is then divided between UCLN and an autocorrelated clock defined on the same set of branch-specific rates. This allows the three familiar clock models to be compared within a single posterior analysis rather than in separate runs followed by external model comparison. In the implementation used here, the top-level strict-versus-relaxed comparison is evaluated analytically, whereas the UCLN-versus-autocorrelated comparison is represented by a binary indicator on the shared relaxed-clock rate vector.

### 2.2 Branch-rate parameterisation

Let *D* denote the alignment, *T* a rooted binary time tree, and *B* the set of non-root branches. For each branch *b* ∈ *B*, let *t*_*b*_ denote its duration in time units and let *λ*_*b*_ denote its substitution rate. The product *t*_*b*_*λ*_*b*_ is the expected evolutionary distance on branch *b* and is the quantity that enters the phylogenetic likelihood. Under the strict clock, all branches share a single rate, so that

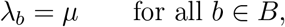

where *µ >* 0 is the overall clock rate. Under the relaxed class, rates are written as

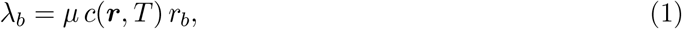

where ***r*** = (*r*_*b*_)_*b*∈*B*_ is a positive vector of branch-specific relative-rate multipliers. We use a real-valued branch-rate parameterisation for UCLN rather than the original discrete-category implementation. Efficient inference under this parameterisation relies on relaxed-clock proposals developed by Zhang and Drummond (2020) and subsequently improved by Douglas et al. (2021). Here the relaxed class contains the two relaxed-clock families considered in this paper: UCLN and the auto-correlated clock. This separates the overall timescale from branch-specific rate variation and allows the same branch-rate vector to be used across the relaxed-clock submodels (Drummond et al., 2006; Kishino et al., 2001; Douglas et al., 2021).

When time-weighted normalisation is used,

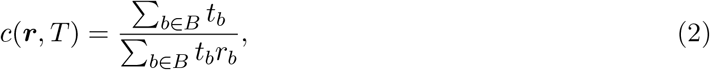

so that *µ* retains the interpretation of the branch-time-weighted mean substitution rate across the tree. Without normalisation, *c*(***r***, *T*) = 1. In the analyses reported here, time-weighted normalisation was enabled. This choice keeps *µ* directly interpretable as the branch-time-weighted mean substitution rate and reduces confounding between *µ* and the overall scale of ***r*** (Drummond et al., 2006; Douglas et al., 2021). Computationally, this normalisation introduces a global dependence: changing a single *r*_*b*_ alters the normalising constant for all branches. We accept that cost here because it keeps *µ* interpretable as the global substitution rate and prevents arbitrary drift between *µ* and the average branch-rate multiplier.

This common parameterisation is used for both relaxed-clock submodels. The uncorrelated relaxed clock treats branch-specific rates as independent draws from a common distribution, whereas the autocorrelated relaxed clock models rates as evolving along the phylogeny so that neighbouring branches tend to have similar values (Drummond et al., 2006; Kishino et al., 2001). The two relaxed clocks therefore differ in the prior process placed on ***r***, not in the definition of the latent branch-rate vector itself. This is the key reason why the within-relaxed comparison can be handled as a fixed-dimensional model-selection problem. This construction differs from the random local clock (Drummond and Suchard, 2010), which assigns rates to phylogenetically contiguous regimes by placing discrete rate-shift indicators on branches; branches that do not carry a shift inherit the rate of their parent, so the number of free rate parameters is itself a random variable. The comparison between uncorrelated and autocorrelated relaxed clocks is therefore fundamentally different in nature: both models define the same full branch-rate vector ***r*** of fixed dimension, and the choice between them reduces to a fixed-dimensional prior-process comparison rather than a trans-dimensional problem over alternative rate-shift configurations.

### 2.3 Top-level mixture over strict and relaxed clocks

For notational convenience, let *ϕ* denote all remaining phylogenetic parameters that are not specific to the clock model. Given a dated tree *T* , an overall substitution rate *µ*, and the observed alignment *D*, let *L*_S_(*D* | *T, µ, ϕ*) denote the phylogenetic likelihood under a strict clock, and let *L*_R_(*D* | *T, µ*, ***r***, *ϕ*) denote the corresponding likelihood under the relaxed part of the model, where ***r*** is the shared branch-rate vector introduced above. We then define the top level of the hierarchical clock model as a two-component mixture,

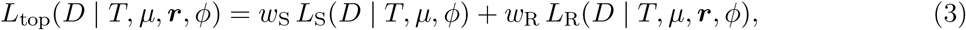

where *w*_S_, *w*_R_ ≥ 0 and *w*_S_ + *w*_R_ = 1. Unlike analytical mixtures that place competing priors on a shared branch-rate vector (Darlim and Höhna, 2024), Equation 3 mixes two likelihoods defined on different parameter spaces: *L*_S_ does not depend on ***r*** at all, whereas *L*_R_ does. The strict component therefore never requires ***r*** to collapse to **1** under a relaxed-clock prior; ***r*** is simply irrelevant to *L*_S_. This is why the top-level mixture can include a strict clock even though prior-level mixtures cannot. In the analyses reported here, we fixed these weights rather than estimating them from the same data. When the inner relaxed-clock weights were set to 1*/*2 and 1*/*2, we used *w*_S_ = 1*/*3 and *w*_R_ = 2*/*3 so that the strict clock, UCLN, and autocorrelated clock each received equal prior mass.

This top-level comparison is evaluated analytically rather than through a sampled allocation variable. At each MCMC state, the strict and relaxed components are combined directly through Equation 3, and the log mixture likelihood is computed with a standard log-sum-exp calculation for numerical stability. The corresponding conditional probabilities are

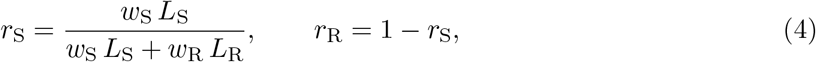

with *r*_S_ + *r*_R_ = 1. If *I* ∈ *{*S, R*}* denotes a latent top-level indicator with prior weights (*w*_S_, *w*_R_), then *r*_S_ and *r*_R_ are Pr(*I* = S | *D, T, µ*, ***r***, *ϕ*) and Pr(*I* = R | *D, T, µ*, ***r***, *ϕ*), respectively. The indicator is marginalised by construction: summing *w*_S_*L*_S_ + *w*_R_*L*_R_ is the marginal likelihood of the two-component model, so the sampler never needs to represent *I* as a state variable. The marginal posterior model probabilities are recovered from these conditional probabilities by averaging over the posterior sample,

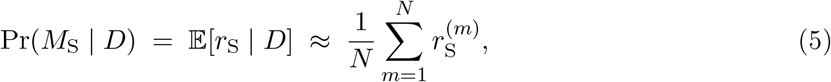

where *N* is the number of retained posterior samples after burn-in, and the approximation improves as *N* increases. The identity follows from the tower property, and similarly for Pr(*M*_R_ | *D*). This Rao–Blackwellized estimator is available at every iteration and avoids the slow mixing that a sampled strict-versus-relaxed indicator would incur when *L*_S_ and *L*_R_ are well separated.

### 2.4 Within-relaxed comparison between UCLN and autocorrelated clocks

Within the relaxed part of the model, we introduce a binary indicator *z* to distinguish between the two alternative descriptions of among-branch rate variation. We write *z* = 0 for UCLN and *z* = 1 for the autocorrelated clock. Conditional on the dated tree *T* , the overall rate *µ*, and the shared branch-rate vector ***r***, the relaxed-clock likelihood is the same under both models. What changes is the prior assigned to ***r***. The within-relaxed part of the hierarchy can therefore be written as

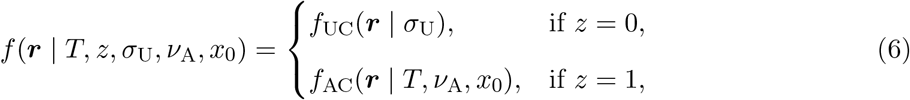

where *σ*_U_ is the UCLN log-scale standard deviation, *ν*_A_ is the autocorrelated diffusion variance per unit time, and *x*_0_ is the root log-rate used by the autocorrelated process.

Under UCLN, the branch-specific relative rates are independent lognormal draws,

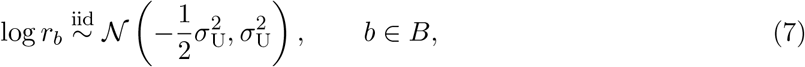

Equivalently, *r*_*b*_ is lognormally distributed with log-scale mean 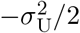 and log-scale variance 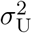, so that E[*r*_*b*_] = 1. Under the autocorrelated clock, log rates evolve along the tree. Writing *x*_*b*_ = log *r*_*b*_, for each non-root branch *b* with parent branch pa(*b*) and duration *t*_*b*_,

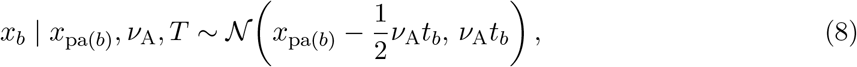

with *x*_pa(*b*)_ = *x*_0_ when the parent node is the root. The mean correction in Equation 8 preserves the rate scale in the sense that E[*r*_*b*_ | *r*_pa(*b*)_] = *r*_pa(*b*)_.

Because the likelihood under the relaxed clock depends on ***r*** but not directly on whether ***r*** is interpreted under UCLN or the autocorrelated prior, the within-relaxed comparison amounts to choosing between two prior models on the same parameter vector. In the analyses reported here, the prior weights for the two relaxed-clock models were fixed at *η*_UC_ = *η*_AC_ = 1*/*2. At any given MCMC state, this gives the conditional posterior probabilities

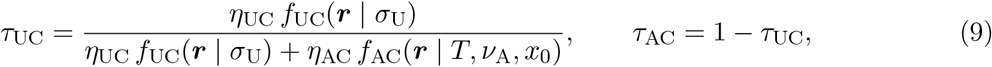

which measure how strongly the current branch-rate configuration favours UCLN or the autocorrelated clock. These quantities are then combined with the top-level strict-versus-relaxed probabilities to obtain the final posterior probabilities for all three clock models.

### 2.5 Posterior probabilities for the three clock models and model-averaged summaries

Combining the top-level probabilities in Equation 4 with the within-relaxed probabilities in Equation 9 gives the posterior probabilities for the three terminal clock models at each MCMC state,

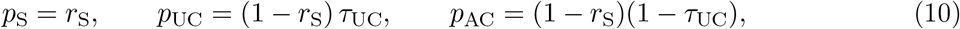

with

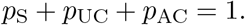

The three terminal probabilities are therefore determined by only two continuous responsibilities (*sensu* Bishop and Nasrabadi, 2006), *r*_S_ and *τ*_UC_, each lying in [0, 1]. They are conditional on the current MCMC state—the current tree, branch rates, and remaining parameters—and so are distinct from the posterior model probabilities, which are their averages over the sample. They are evaluated analytically at every state and are not binary indicators: a single state contributes fractionally to all three clock families rather than being assigned to one of them. The only genuinely sampled *{*0, 1*}* variable in the model is the within-relaxed indicator *z*; the cross-model operators (Section 2.6) update *z* and remap the branch-rate vector between the UCLN and autocorrelated parameterisations, so that ***r*** is explored under both relaxed-clock priors. By the same Rao–Blackwellization argument as in Equation 5, averaging the per-iteration 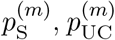, and 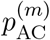 over the posterior sample gives consistent estimates of Pr(*M*_S_ | *D*), Pr(*M*_UC_ | *D*), and Pr(*M*_AC_ | *D*). We report these responsibilities rather than the empirical frequencies of the sampled indicator *z* because, being Rao–Blackwellized, they have lower Monte Carlo variance and remain informative even when *z* mixes slowly.

For quantities that are shared across all three clock models, such as the dated tree, the overall substitution rate, root age, or tree length, model-averaged posterior summaries are obtained directly from the full MCMC output, since the posterior already averages over clock uncertainty. For quantities that are specific to one relaxed-clock model, however, summaries should be reported conditionally on that model. If *θ*_UC_ denotes a parameter defined under UCLN and *θ*_AC_ a parameter defined under the autocorrelated clock, we estimate

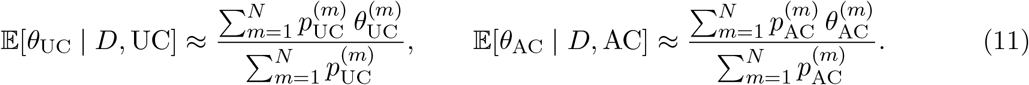

These responsibility-weighted averages define model-specific posterior means conditional on the corresponding relaxed-clock family. They keep the reported summaries aligned with the three-model posterior probabilities in Equation 10, rather than treating every visit to the relaxed part of the model as equally informative about both relaxed-clock models.

Throughout the paper, we therefore distinguish between two kinds of posterior summary. The first is a model-averaged summary for quantities whose meaning does not depend on which clock model is used. The second is a model-specific summary for quantities that only arise under one of the two relaxed-clock models. This distinction is important in practice, because the data may strongly reject a strict clock while still leaving appreciable uncertainty about whether among-branch rate variation is better described by UCLN or by an autocorrelated process.

### 2.6 MCMC proposals for the clock mixture

The within-relaxed indicator *z* ∈ *{*0, 1*}* is the non-trivial mixing problem in this model. A naive Metropolis–Hastings flip of *z* without coherent adjustment of the branch-rate vector ***r*** almost always proposes a rate configuration that is implausible under the proposed model’s prior, so the chain becomes stuck on whichever clock family it started in. We therefore used two specialised cross-model operators for *z*: an exact Gibbs update (`IndicatorGibbsOperator`) that draws *z* from its full conditional given ***r*** and the hyperparameters, and a deterministic quantile-preserving bridge (`UCACSwitchBridgeOperator`) that flips *z* while coherently remapping ***r*** between the UCLN and autocorrelated non-centered parameterisations. The Gibbs update is standard Bayesian variable selection (George and McCulloch, 1993; Kuo and Mallick, 1998) and accepts with probability one. The deterministic bridge extends the quantile-parameterisation idea of Li and Drummond (2012) to cross-model moves between UCLN and the autocorrelated clock, with the Jacobian required when the MCMC state is the branch-rate vector itself.

Within-component mixing of ***r*** under the autocorrelated model is improved by a block proposal (`ACSubtreeUIncrementOperator`) that perturbs the whitened residuals of a random subtree under the non-centered parameterisation (Papaspiliopoulos et al., 2007) and pushes them back through the autocorrelated map. Non-centered scale moves on the relaxed-clock hyperparameters *σ*_U_ and *ν*_A_ (`UCLDStdevNonCenteredOperator` and `ACSigma2NonCenteredOperator`) co-transform ***r*** in lockstep with the hyperparameter so that each branch-rate retains its quantile position under the relaxed-clock prior.

The UC–AC bridge showed data-dependent behaviour, with acceptance ranging from ∼ 10^−5^ on the 31-taxon *rbcL* JTT analysis to ∼ 0.5 on DENV-4; we therefore retained it as a complementary long-range proposal alongside the Gibbs update. Full derivations, Jacobians, operator weights, and per-analysis acceptance diagnostics are given in Supplementary Note 4.

### 2.7 Stratified simulation study

We conducted a stratified simulation study under correct model specification by generating 100 replicate alignments under each of the three terminal clock models, so that performance could be assessed conditional on the generating clock family. Using LPhyStudio and LPhyBEAST within the LinguaPhylo framework (Drummond et al., 2023), we simulated 100 replicate alignments under a strict clock, 100 under UCLN, and 100 under the autocorrelated clock, giving 300 data sets in total.

Across all three strata, 30 heterochronously sampled taxa were placed at one-third-unit intervals, genealogies were drawn from a constant-size coalescent with Θ ∼ LogNormal(1.2, 0.6) (Kingman, 1982a,b), and alignments of 1500 nucleotides were simulated under JC69 with four-category discrete-gamma site-rate variation, with *γ* ∼ LogNormal(−1.0, 0.7) (Jukes et al., 1969; Yang, 1994). JC69 was chosen deliberately to isolate clock-model behaviour from substitution-model misspecification; the aim of the simulation study was to evaluate recovery of clock-family support under a controlled and minimal substitution process rather than to mimic the full complexity of empirical data. The overall substitution rate was drawn as *µ* ∼ LogNormal(−3.4, 0.5) in all three strata. We also recorded the mean-rate-scaled root age *µH*, where *H* is the true root age in time units. The median *µH* was 0.50 substitutions site^−1^ (central 95% range 0.17–1.70). For strict-clock simulations, all branches shared this single rate. For UCLN simulations, the log-scale standard deviation was drawn as *σ*_U_ ∼ LogNormal(−1.1, 0.6). For autocorrelated simulations, the variance parameter was drawn as *ν*_A_ ∼ LogNormal(−4, 1.0), with the root log-rate fixed at *x*_0_ = 0. Each replicate was then analysed under the full hierarchical mixture model using the same priors, fixed mixture weights, and operator settings as in the main analyses. Full simulation scripts are provided in the Supplementary Material. As an additional check outside this regime, we ran a further set of simulations under a lower substitution rate (*µ* ∼ LogNormal(−5, 0.5)) and shorter alignments (600 nucleotides), placing the data in a lower-information regime (median *µH* = 0.10 substitutions site^−1^, central 95% range 0.04–0.29). Results are reported in the Supplementary Material.

The simulation study addressed two questions. First, we asked whether the analysis placed posterior support on the correct generating clock model. For replicate *j*, let 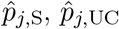, and 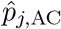 denote the posterior means of Equation 10. We summarised discrete-model recovery by whether the true generating model was contained in the 95% posterior model set, defined as the smallest set of clock models whose cumulative posterior probability reached the nominal 95% level.

Second, we asked whether carrying clock uncertainty through the analysis still allowed accurate estimation of quantities shared across the three clock models. For each replicate, we recorded the posterior median and 95% highest posterior density interval for the overall substitution rate, root age, and total tree length, and compared these summaries with the true generating values. Recovery was assessed by log-scale estimation error and empirical 95% interval coverage. Because these quantities are shared across clock models, all summaries were taken from the full model-averaged posterior rather than from model-specific conditional distributions.

### 2.8 Empirical analyses

The four empirical data sets benchmarked different aspects of the method: DENV-4 against standalone evidence-based model comparison, the 31-taxon *rbcL* set of Thorne et al. (1998) as a historical benchmark for autocorrelated-rate inference, and RSV-A and H3N2 as contemporary viral applications differing in scale and phylogenetic context. All mixture analyses used the same prior masses as in the simulation study so that strict, UCLN, and autocorrelated each received prior probability 1*/*3, with non-clock model components held fixed across clock-model comparisons.

For DENV-4, we analysed the 17-taxon serially sampled envelope-gene alignment (1485 bp) spanning 1956–1994 compiled by Rambaut (2000) from the Dengue-4 sequences of Lanciotti et al. (1997), with one recombinant sequence identified by Worobey et al. (1999) omitted. We used an HKY site model with estimated base frequencies, four-category discrete-gamma rate heterogeneity, and a constant-size coalescent tree prior. We additionally fitted standalone strict-clock, UCLN, and autocorrelated analyses under the same non-clock model and estimated their marginal likelihoods by nested sampling. To assess practical stability, we used both an exploratory configuration and a higher-precision configuration, and for each configuration we repeated the three standalone analyses across replicate triplets. Nested-sampling log marginal likelihoods were then converted to posterior model probabilities under equal prior model weights, allowing direct comparison with the posterior probabilities returned by the mixture analysis.

The historical benchmark was the 31-taxon chloroplast *rbcL* amino-acid data set used by Thorne et al. (1998) to illustrate autocorrelated rate evolution, reconstructed from GenBank accessions associated with the original *rbcL* sampling of Chase et al. (1993). *Marchantia paleacea* was retained as the outgroup following Thorne et al. (1998). The baseline mixture analysis used a JTT amino-acid replacement model without gamma-distributed rate heterogeneity or an invariant-sites component, together with a Yule tree prior and a monophyly constraint on the 30 ingroup taxa. As a site-model sensitivity analysis, we repeated the mixture under OBAMA (Bouckaert, 2020), a BEAST 2 framework for Bayesian amino-acid model averaging, keeping the same tree prior and clock-mixture settings.

The BEAST RSV2 benchmark is a serially sampled RSV-A G-gene alignment of 129 sequences (629 nt) collected between 1956 and 2002, comprising 81 Belgian isolates and 48 previously published sequences retrieved from GenBank (Table 1 of Zlateva et al. (2004)). It was analysed without sub-sampling or other custom preprocessing, under an HKY site model with estimated base frequencies and a constant-size coalescent tree prior. For this data set, we recorded posterior probabilities for the three clock families together with the overall substitution rate and root age from the full model-averaged posterior, and the UCLN standard deviation as a UCLN-specific summary, so that the mixture analysis could be compared directly with the classical RSV-A timescale reported by Zlateva et al. (2004).

**Table 1:** Summary of simulation-based validation under the three generating clock models.

| True model | In 95% set (%) | Credible set size (%) |  |  | HPD coverage (%) |  |  |
| --- | --- | --- | --- | --- | --- | --- | --- |
|  |  | Size 1 | Size 2 | Size 3 | Clock rate | Root age | Tree length |
| Strict | 100 | 36 | 64 | 0 | 96 | 93 | 93 |
| UCLN | 100 | 95 | 5 | 0 | 96 | 95 | 92 |
| Autocorrelated | 100 | 82 | 18 | 0 | 95 | 93 | 92 |

The H3N2 structured-coalescent benchmark of Vaughan et al. (2014) comprises 980 HA sequences from Hong Kong, New York, and New Zealand sampled between 2000 and 2006. Because exact structured-coalescent inference at the full 980-taxon scale was computationally demanding, we analysed a stratified quarter-sized subset of 245 sequences (55 Hong Kong, 79 New York, 111 New Zealand; sampling times 2000.005–2006.175), drawn by proportional largest-remainder sampling across region *×* calendar-year strata. The sequence model was GTR with estimated base frequencies and four-category discrete-gamma rate heterogeneity, and the tree prior was the exact three-deme structured coalescent with lognormal priors on deme sizes and the backward migration-rate matrix.

For H3N2, we recorded posterior probabilities for the three clock families and model-averaged summaries for the overall substitution rate, root age, and the three deme-size parameters. We used the published 980-taxon strict-clock structured-coalescent analysis of Vaughan et al. (2014) as an external reference for the inferred timescale and broad structured-coalescent summaries, rather than as a same-data model-comparison control. For consistency with that reference, H3N2 summaries are reported as posterior medians and 95% highest posterior density intervals. For branch-level visualisation, we generated a CCD0-MAP summary tree (Berling et al., 2025) for the RSV-A analysis and coloured branches by the posterior branch-rate summaries logged by the hierarchical mixture analysis. For the rbcL OBAMA sensitivity analysis, we compared the two relaxed-clock families directly by computing UCLN-conditioned and autocorrelated-clock-conditioned posterior mean branch-rate multipliers for each branch. Branches were identified across posterior trees by their descendant taxon sets, and conditional means were weighted by the terminal responsibilities *p*_UC_ and *p*_AC_ from Equation 10. For the branch-wise comparison, we retained only branches with posterior frequency at least 0.5 in the posterior tree sample. Analogous branch-level diagnostics were considered for all empirical analyses whose posterior tree logs contained branch-rate metadata. Supplementary Note 5 summarises which diagnostics were displayed, archived, or not interpretable from the available tree logs. We interpret UCLN-versus-autocorrelated branch-rate comparisons only when both relaxed-clock families retain non-negligible posterior support; otherwise the corresponding plots are treated as diagnostics rather than primary branch-level summaries.

### 2.9 Software implementation

The method is implemented in the open-source `RelaxClockAveraging` package for BEAST 2. The package has four main modelling components: a top-level mixture likelihood for strict-versus-relaxed comparison, a shared-rates clock model for the relaxed class, a stochastic-variable-selection prior for the within-relaxed boundary, and logging utilities for reconstructing posterior family probabilities and conditional summaries. Routine within-model exploration of the shared rate vector uses standard multiplicative scale proposals on individual rates and on subtree blocks, together with the non-centered autocorrelated subtree proposal and non-centered hyperparameter proposals summarised above (Section 2.6; full derivations in Supplementary Note 4).

Operators whose proposal is meaningful only under one value of the indicator, including these non-centered operators, are all registered in a single standard BEAST 2 operator schedule and are selected at fixed, state-independent rates. When an operator is selected in an indicator state in which it has no valid proposal, it performs no state change and returns a log Hastings ratio of −∞, so the draw is rejected at no further cost. This design costs a small amount of efficiency, because operators are occasionally selected in states where they cannot act, but it keeps the operator-selection probability state-independent, so the standard Metropolis–Hastings acceptance ratio applies to every operator without correction. A state-conditional operator schedule that selected only currently-applicable operators would reclaim the lost efficiency, but every operator that changes the indicator would then require an additional log(*W*_pre_*/W*_post_) correction in its Hastings ratio to compensate for the difference in operator-selection probability before and after the move, where *W*_pre_ and *W*_post_ are the total schedule weights active in the pre- and post-move states. This correction would silently change any time an unrelated operator was added to or removed from either sub-schedule, a failure mode that is easy to overlook. For a study whose primary output is a posterior distribution over clock-family indicators, we judged the fixed-schedule design to be the safer choice.

## 3 Results

### 3.1 Inference from simulated data

Across the 300 simulated data sets, the hierarchical analysis consistently retained the true generating clock model within the 95% posterior model set (Fig. 1; Table 1). Coverage was 100% for all three clock families—strict-clock, UCLN, and autocorrelated simulations alike.

**Figure 1.**
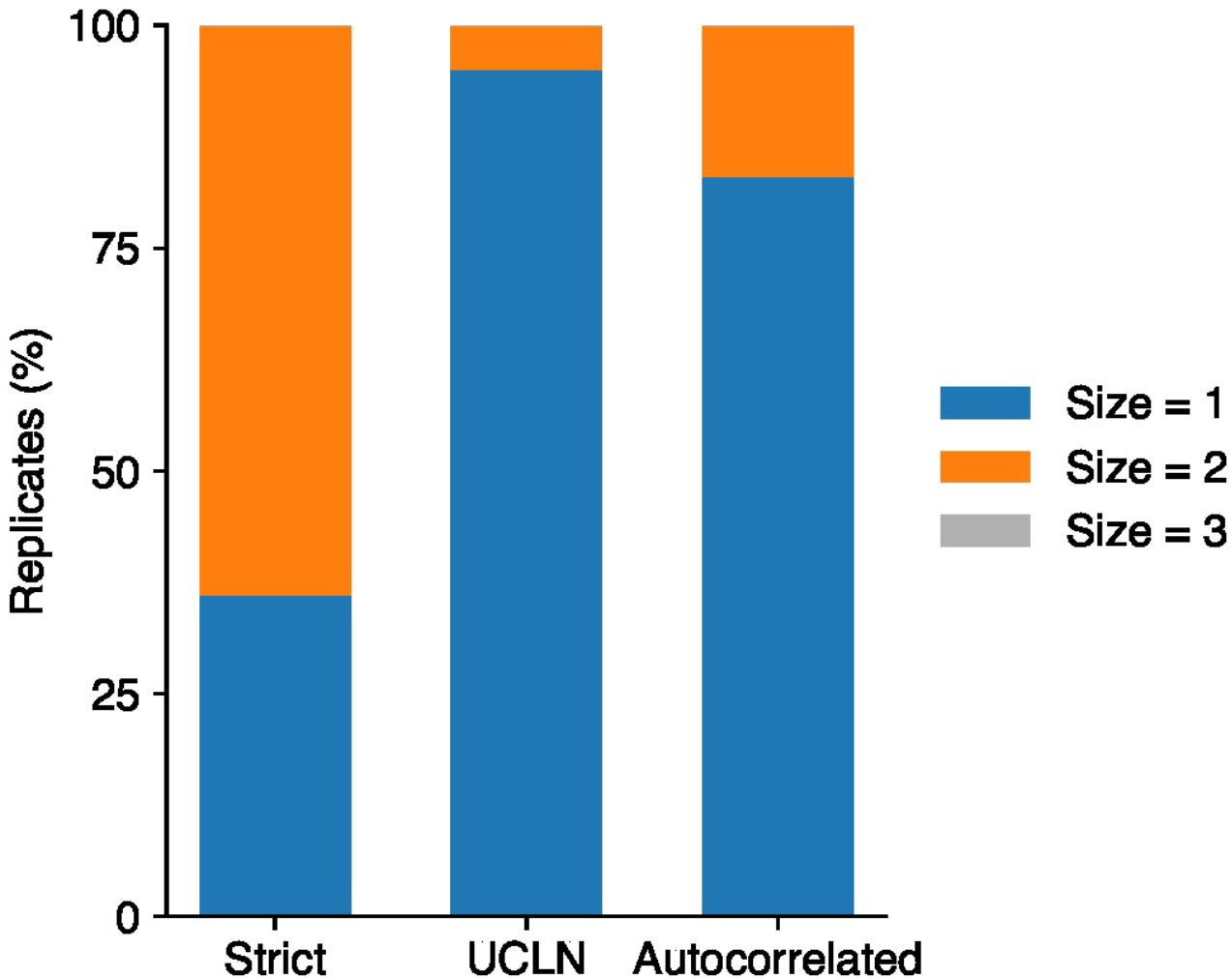
Stratified simulation study of the size of the 95% posterior model set across 300 replicates. Bars show the percentage of replicates for which the 95% posterior model set contained one, two, or three clock models, stratified by generating clock family. Most replicates yielded a singleton model set, while a minority retained two clock models and none retained all three. The true generating clock model was contained in the 95% posterior model set in all replicates; exact coverage values are reported in Table 1.

These coverage values, however, should be interpreted together with the size of the posterior model set. Across the 300 replicates, 213 yielded a singleton 95% model set, whereas the remaining 87 retained two clock models; no replicate required all three. The posterior was most diffuse for strict-clock simulations, for which the 95% model set contained more than one clock model in 64 of 100 replicates. Singleton sets were considerably more frequent when the true generating model was UCLN or autocorrelated (95 and 82 of 100 replicates, respectively), but even under these two relaxed-clock conditions the posterior occasionally retained a second plausible clock model. Under this simulation design, the method therefore behaves conservatively at the model level: it did not exclude the true generating clock model in any replicate, but it sometimes carried clock-model uncertainty forward rather than collapsing to a single clock family.

Despite this residual model-level uncertainty, model-averaged estimation of the shared continuous parameters remained accurate. For the overall substitution rate, empirical 95% highest posterior density interval coverage was 96%, 96%, and 95% under strict-clock, UCLN, and autocorrelated simulations, respectively. The corresponding values were 93%, 95%, and 93% for root age, and 93%, 92%, and 92% for total tree length.

These results show that the hierarchical analysis retains the true clock model within the posterior model set at or above nominal levels, preserves uncertainty when the three clock families are not sharply separated by the data, and still yields accurate model-averaged estimation of shared timescale parameters.

### 3.2 Nested-sampling benchmark on the DENV-4 data

We compared the three clock families on the DENV-4 benchmark using both the single-run hierarchical mixture and standalone nested-sampling analyses for each clock model. At the exploratory nested-sampling setting of 30 particles and a subchain length of 5,000, nested sampling was highly unstable. Replicate-to-replicate variation in log marginal likelihood was broad for all three clock models, with substantial overlap among their empirical distributions (Fig. 2B). The strict clock ranked first in only 9 of 20 replicate triplets, with the UCLN and autocorrelated clocks ranking first in 7 and 4 triplets, respectively, and the preferred model changed repeatedly across replicates. The NS uncertainty reported internally by BEAST was appreciably smaller than the observed replicate-to-replicate dispersion, indicating that the 30/5,000 configuration is not sufficiently stable for definitive model comparison on this data set.

**Figure 2.**
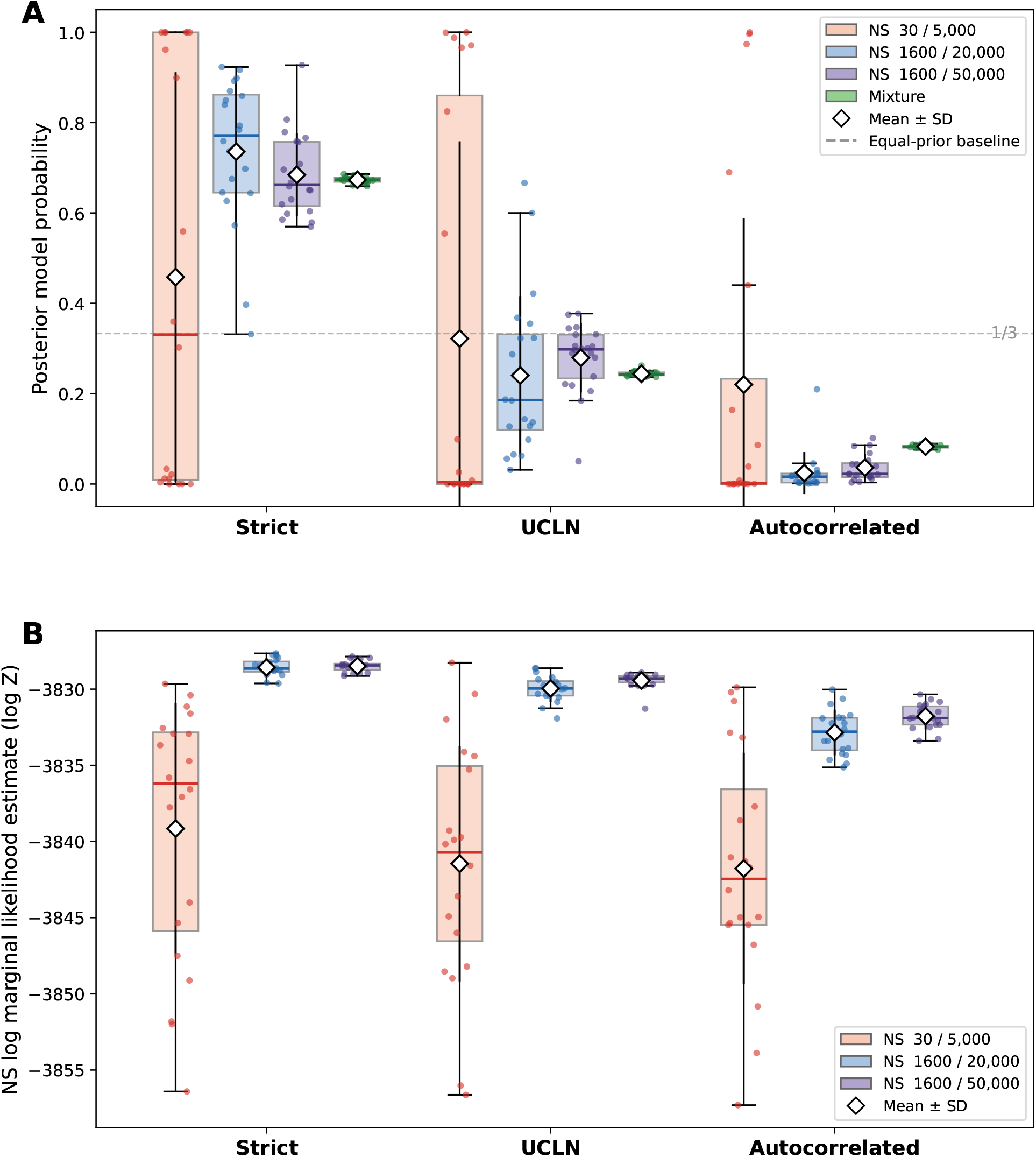
Nested-sampling benchmark on the DENV-4 data. (A) Posterior model probabilities obtained by converting replicate-wise log marginal likelihoods under equal prior model masses; the mixture posterior is shown on the same three-model probability scale for direct comparison. (B) Replicate variation in NS log marginal likelihood across three computational settings. Boxes show interquartile ranges, centre lines show medians, whiskers extend to the most extreme observations within 1.5 interquartile ranges, small points denote individual replicate runs or triplets, and diamonds denote mean *±* 1 standard deviation across replicates. The low-cost NS configuration is highly unstable, whereas the higher-effort settings stabilise the qualitative ranking strict *>* UCLN *>* autocorrelated. The dashed line in panel (A) marks the equal-prior baseline of 1*/*3.

Stabilising the NS benchmark required an increase in computational effort of roughly two orders of magnitude: from 30 particles *×* 5,000 subchain to 1,600 particles *×* 20,000 subchain, a roughly 200-fold increase in nested-sampling work relative to the exploratory setting. At that cost, the replicate standard deviation of the log marginal likelihood fell to 0.554 for the strict clock, 0.835 for the UCLN clock, and 1.466 for the autocorrelated clock, and the strict clock ranked first in 18 of 20 replicate triplets. A further 1,600/50,000 configuration produced the same overall ordering and similarly tight log-marginal-likelihood distributions, confirming that the qualitative ranking strict *>* UCLN *>* autocorrelated only becomes reliable once NS is run at this level of effort. Even then, the two higher-effort NS settings did not agree on a single posterior allocation across the three clock models, showing that stabilising the model ranking is not the same as accurately estimating the posterior model probabilities.

This distinction is clearest on the posterior probability scale (Fig. 2A). Under the 30/5,000 configuration, posterior support remained diffuse across the three clock models. Under both higher-effort NS configurations, posterior mass concentrated on the strict clock, the UCLN clock retained secondary support, and the autocorrelated clock received little support. However, the exact posterior mass assigned to the strict and UCLN clocks remained somewhat sensitive to the NS configuration even after the log-marginal-likelihood ranking had stabilised. The highest-effort NS configuration (1,600/50,000) did not produce posterior probabilities systematically closer to the mixture result than the 1,600/20,000 setting.

The computational contrast was also substantial. BEAST-reported calculation times were summarised from runs per clock model and setting on eight cores per run. The mixture analysis required a median of 2.78 wall-clock hours (range 2.77–3.50 h) to reach the reported posterior summaries; in the run used for those summaries, all primary clock- and tree-related trace effective sample sizes were above 900. By contrast, a complete three-clock nested-sampling comparison required three separate standalone analyses. The low-cost 30/5,000 nested-sampling setting was inexpensive, with a median three-clock triplet work of 1.05 run-hours (model-wise summed range 0.77–1.70 h), but Fig. 2 shows that it was not stable enough for reliable model comparison. The relevant computational contrast is therefore with the higher-effort settings that stabilised the model ranking. For those settings, a complete three-clock nested-sampling comparison required 198.5 run-hours (model-wise summed range 179.6–305.5 h) and 675.4 run-hours (452.4–716.8 h) under the 1,600/20,000 and 1,600/50,000 configurations, respectively, compared with 2.78 h for the mixture analysis. Using the sum of the median strict, UCLN, and autocorrelated runtimes as a measure of computational work, these correspond to approximately 1,588 and 5,403 core-hours, or about 71-fold and 243-fold more computational work than a single mixture analysis. Because the three standalone clock analyses can be launched concurrently, these sums should not be interpreted as minimum calendar waiting times; even under full parallelisation across the three clock models, however, the elapsed time would be governed by the slowest run, with median slowest-model times of 85.1 h (range 82.6–135.1 h) and 285.2 h (176.9–299.6 h) for the 1,600/20,000 and 1,600/50,000 settings. Thus the higher-effort nested-sampling configurations that stabilised the model ranking were substantially more expensive than the single joint mixture analysis.

The mixture analysis agreed qualitatively with the higher-effort NS benchmarks by placing the strongest support on the strict clock, weaker but non-zero support on the UCLN clock, and low support on the autocorrelated clock. The mixture does not remove posterior uncertainty. It estimates support for all three clock models within a single joint analysis, avoiding the additional run-to-run variability that arises when posterior model probabilities are assembled from separately estimated marginal likelihoods. The observation that this level of computation was required on a data set of only 17 taxa underscores the practical burden of obtaining stable clock-family posterior model probabilities from repeated standalone evidence calculations, and helps motivate the single-analysis alternative developed here.

### 3.3 The 31-taxon *rbcL* benchmark for autocorrelated-rate inference

Under the baseline JTT amino-acid analysis, the 31-taxon chloroplast *rbcL* benchmark showed strong support for the autocorrelated clock. Posterior probability was concentrated almost entirely on the autocorrelated model (*P*_AC_ = 0.968), with only minor support for UCLN (*P*_UC_ = 0.032). This result is important less as a new biological claim than as a historical check on the method: the data set was originally used to illustrate autocorrelated rate evolution, and the present analysis shows that allowing the strict, UCLN, and autocorrelated clocks to compete within a single posterior framework retains that classical signal.

As a site-model sensitivity analysis, we repeated the benchmark under OBAMA, a Bayesian amino-acid site-model averaging framework. In that analysis, the strict clock remained effectively excluded (*P*_S_ ≈ 3.8 *×* 10^−4^), while posterior support within the relaxed class was divided between UCLN (*P*_UC_ ≈ 0.409) and the autocorrelated clock (*P*_AC_ ≈ 0.591), with a modest preference for the autocorrelated family. We interpret this shift as evidence that the historical benchmark strongly disfavours a strict clock, whereas the internal distinction between UCLN and autocorrelated relaxed clocks is sensitive to the amino-acid site model. The OBAMA posterior placed effectively all substitution-model mass on JTT, and strongly supported both gamma-distributed site-rate heterogeneity and an invariant-sites component. This richer site model allowed part of the heterogeneity that the fixed JTT analysis had to attribute to lineage-specific rate structure to be absorbed at the site level. Root-age summaries changed comparatively little across the two analyses, whereas total tree length was more sensitive, consistent with a redistribution of branch-length allocation rather than a complete change in the overall temporal depth of the tree.

Because the OBAMA analysis assigned substantial posterior support to both relaxed-clock families while assigning negligible support to the strict clock, it also provides a useful branch-level comparison between the two relaxed rate processes. UCLN-conditioned and autocorrelated-clock-conditioned posterior mean branch-rate multipliers were broadly concordant across retained branches (Fig. 3), although some branches showed moderate departures from equality. Thus, in this sensitivity analysis, changing the site model primarily altered posterior allocation between relaxed-clock families rather than producing radically different branch-wise rate summaries.

**Figure 3.**
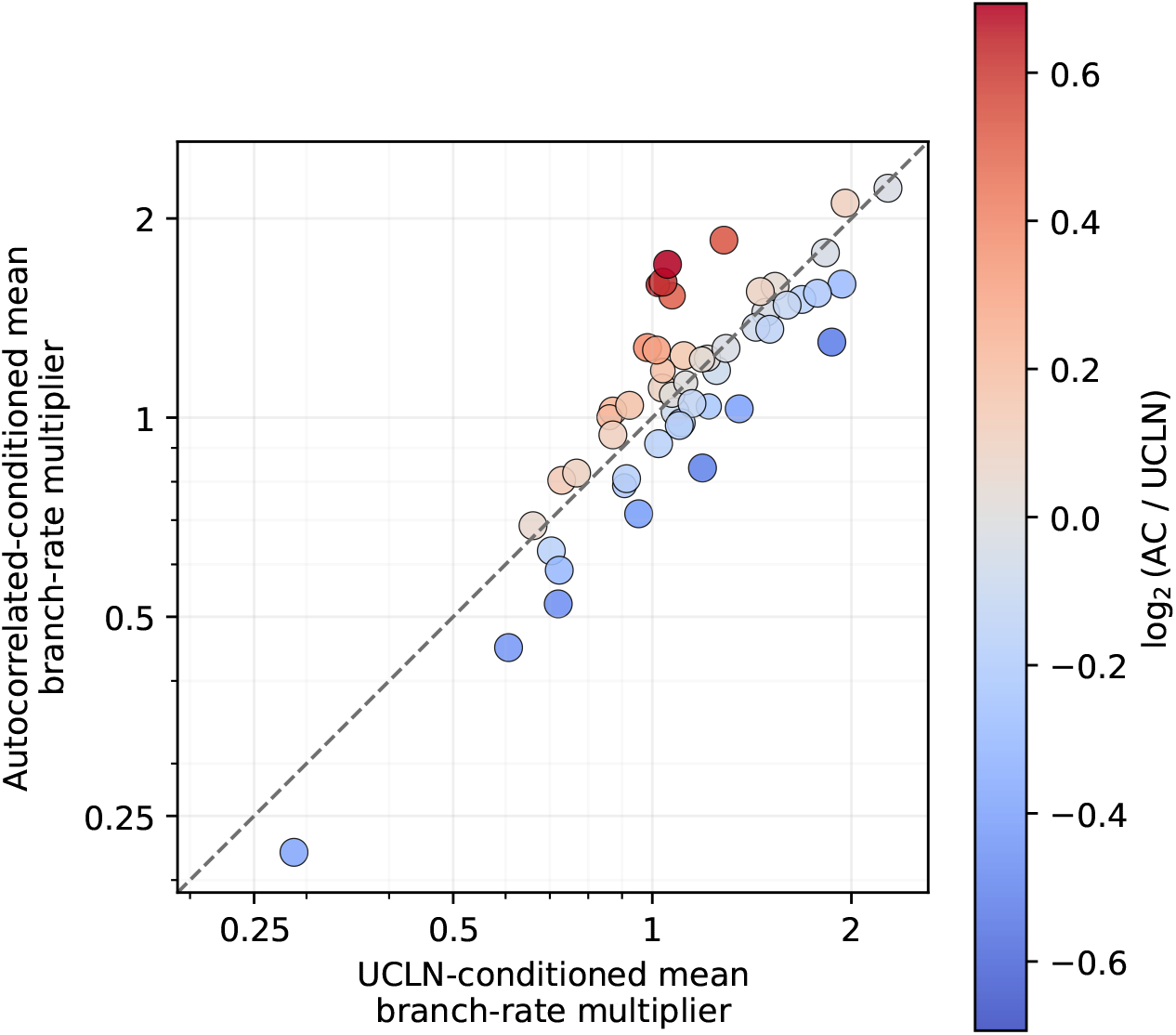
Branch-wise comparison of UCLN-conditioned and autocorrelated-clock-conditioned posterior mean branch-rate multipliers for the *rbcL* OBAMA sensitivity analysis. Each point corresponds to a branch with posterior frequency at least 0.5 in the posterior tree sample. Point size indicates branch posterior frequency, and point colour indicates log_2_(AC*/*UCLN), where zero corresponds to equal conditional branch-rate multipliers under the two relaxed-clock families. The dashed line marks equality between the two conditional branch-rate estimates. Branch-rate values are relative multipliers rather than absolute substitution rates.

### 3.4 The RSV-A G-gene benchmark

On the RSV-A benchmark, posterior mass concentrated almost entirely on the UCLN family (*P*_UC_ ≈ 0.991), with minor residual support for the autocorrelated clock (*P*_AC_ ≈ 9.2 *×* 10^−3^) and effectively none for the strict clock (*P*_S_ ≈ 2.1 *×* 10^−9^). Given this near-degenerate allocation, model-averaged summaries were numerically indistinguishable from UCLN-conditional summaries. The posterior mean UCLN standard deviation was 0.456 (95% HPD 0.30–0.61), indicating substantial among-branch rate variation. The CCD0-MAP summary tree provides a branch-level view of this heterogeneity, with branches coloured by posterior branch-rate summaries from the hierarchical mixture analysis (Fig. 4). The model-averaged posterior mean overall substitution rate was 2.34 *×* 10^−3^ substitutions site^−1^ year^−1^ (95% HPD (2.01–2.70) *×* 10^−3^), and the posterior mean root age was 58.8 years (95% HPD 49.7–69.2), implying a tMRCA of 1943.2 (95% HPD 1932.8–1952.3) relative to the most recent sample year.

**Figure 4.**
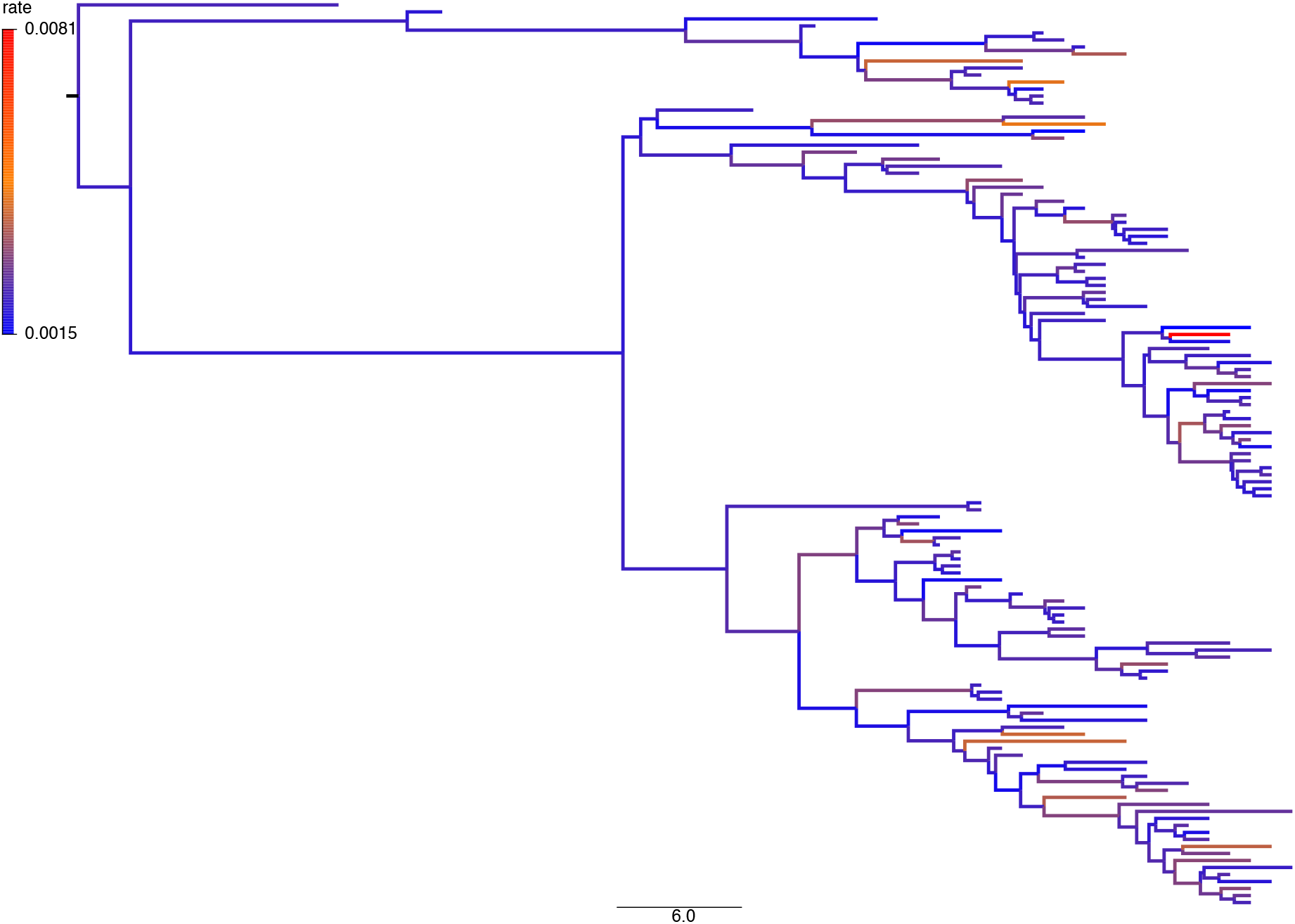
CCD0-MAP summary tree for the RSV-A G-gene benchmark. Each branch is coloured by its posterior branch-rate summary from the hierarchical clock-mixture analysis. Colour-bar values are in substitutions site^−1^ year^−1^, and branch lengths are in years. Because posterior support for RSV-A was concentrated almost entirely on UCLN (*P*_UC_ ≈ 0.991), with only minor autocorrelated support (*P*_AC_ ≈ 9.2 *×* 10^−3^) and negligible strict-clock support, the displayed branch-rate pattern is effectively UCLN-dominated.

These timescale estimates are close to those reported in the classical analysis of Zlateva et al. (2004). In that study, an exploratory root-to-tip regression yielded an evolutionary rate of 1.47 *×* 10^−3^ substitutions site^−1^ year^−1^ (95% CI (1.11–2.18) *×* 10^−3^) and a tMRCA of 1940 (95% CI 1921–1948), while the SRDT analysis gave 1.83 *×* 10^−3^ substitutions site^−1^ year^−1^ (95% CI (1.44–2.26) *×* 10^−3^) and a tMRCA of 1944 (95% CI 1937–1950). Zlateva et al. also reported that a single-rate molecular clock was rejected by a genealogy-based maximum-likelihood test (*P <* 0.01). We therefore interpret this benchmark as consistent with the historical RSV-A literature, while making the clock-family allocation explicit: the hierarchical mixture preserves the established timescale while translating earlier evidence against a single-rate clock into explicit posterior support for the UCLN family.

### 3.5 Reanalysis of the H3N2 structured-coalescent benchmark

H3N2 was our most computationally demanding empirical application, combining dense serial sampling with a three-deme structured coalescent on a 245-taxon stratified subset of the 980-taxon alignment of Vaughan et al. (2014). All parameters discussed below had effective sample sizes above 200.

Unlike the RSV benchmark, H3N2 did not collapse onto a single clock family. The terminal posterior model probabilities were *P*_S_ = 0.256, *P*_UC_ = 0.128, and *P*_AC_ = 0.615. Thus, the top-level strict-versus-relaxed comparison assigned posterior probability *P*_R_ = *P*_UC_ + *P*_AC_ = 0.743 to the relaxed class, corresponding to relaxed-versus-strict posterior odds of approximately 2.9:1. Because the relaxed class received prior mass 2*/*3 under the equal terminal model weights used here, this represents a moderate shift toward relaxation rather than a decisive exclusion of the strict clock. Conditional on relaxation, however, support was concentrated on the autocorrelated clock: Pr(*M*_AC_ | *D, M*_R_) = *P*_AC_*/*(*P*_UC_ + *P*_AC_) = 0.615*/*0.743 ≈ 0.828. The corresponding autocorrelated-versus-UCLN posterior odds within the relaxed class were approximately 4.8:1. The H3N2 subset therefore provides substantial, but not overwhelming, posterior support for the autocorrelated family: most posterior mass lies in the relaxed class, and most relaxed-class support lies on the autocorrelated clock, but the strict clock retains non-negligible posterior probability.

We report posterior medians and 95% HPD intervals, following the convention of Vaughan et al. (2014), and use their published 980-taxon strict-clock analysis as an external timescale reference rather than a same-data model-comparison control. Model-averaged estimates from the quarter-sized mixture analysis were close to the published full-data summaries (Fig. 5). The model-averaged overall substitution rate was 4.434*×*10^−3^ substitutions site^−1^ year^−1^ (95% HPD (3.82–5.11)*×*10^−3^), compared with 5.03 *×* 10^−3^ (95% HPD (4.58–5.51) *×* 10^−3^) in the published 980-taxon strict-clock analysis. The corresponding root age was 7.766 years (95% HPD 7.16–8.50), close to the published 7.62 years (95% HPD 7.21–8.11). Relative to the most recent sample in 2006.175, this implies a root date of 1998.41 (95% HPD 1997.67–1999.01). The H3N2 samples span 2000.005–2006.175, an interval that is itself a large fraction of the inferred root age of roughly eight years, so the timescale is strongly constrained by the sampling dates (Drummond et al., 2002) and is correspondingly insensitive to clock-model choice; the agreement with the published estimate should be read in that light rather than as evidence that clock choice does not matter for timescale estimation in general.

**Figure 5.**
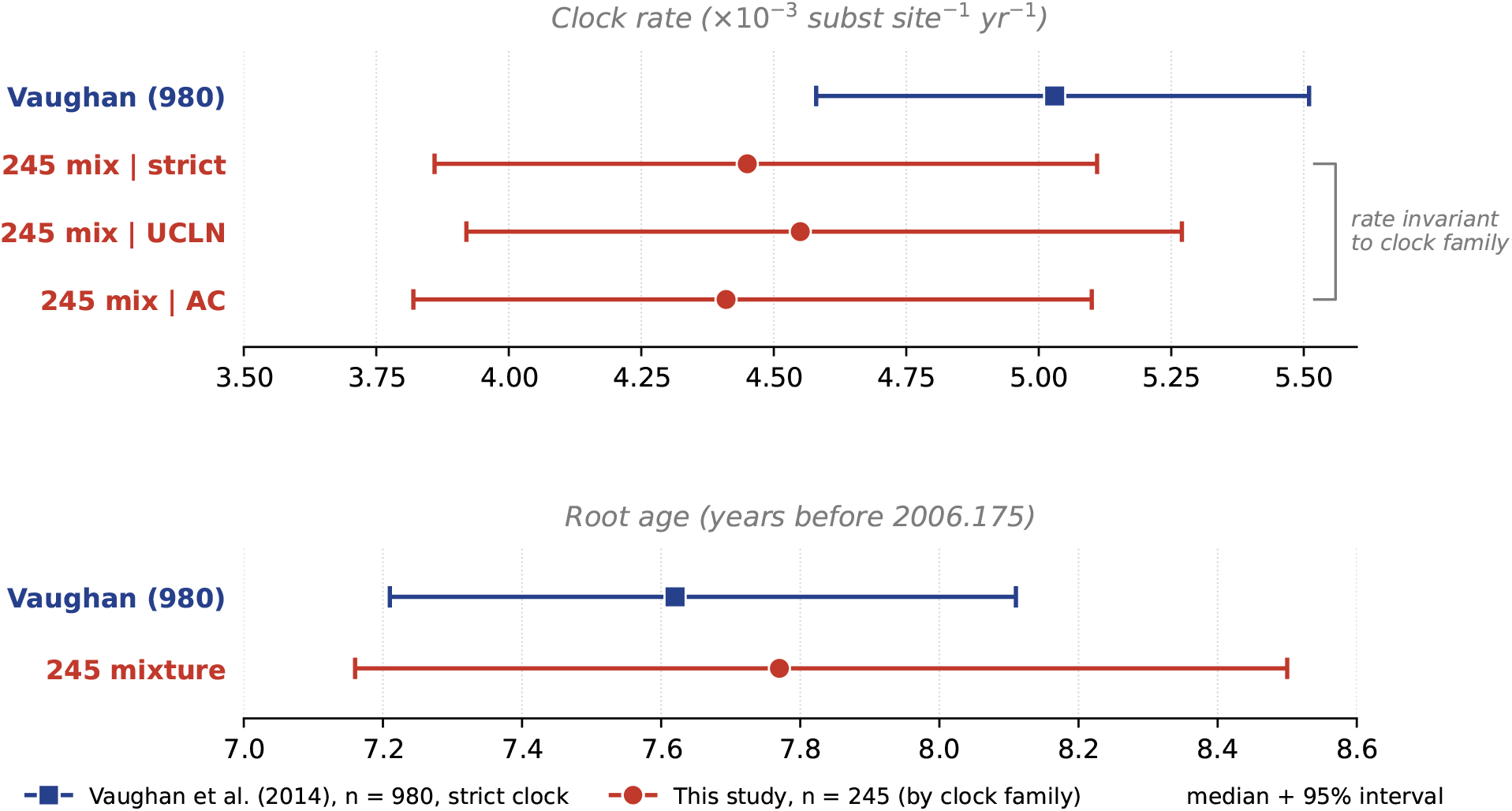
H3N2 timescale under the 245-taxon hierarchical clock mixture, compared with the published 980-taxon strict-clock structured-coalescent analysis of Vaughan et al. (2014). **Top:** the overall substitution rate, reported as the branch-time-weighted tree-wide rate 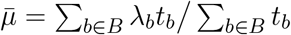 (substitutions site^−1^ year^−1^). For the 245-taxon mixture, 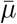 is shown conditional on each clock family as a responsibility-weighted posterior mean, weighting every posterior draw by the per-iteration family responsibilities (Eq. 11): 4.46 (strict), 4.57 (UCLN), and 4.42 (autocorrelated), all *×* 10^−3^. The three conditional estimates overlap almost completely, so 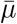 is essentially invariant to clock-family choice; the offset relative to the 980-taxon strict-clock estimate (5.03 [4.58, 5.51] *×* 10^−3^) therefore reflects the difference in taxon sampling between the two analyses rather than clock-model choice. **Bottom:** root age. The 245-taxon mixture gives a model-averaged posterior median of 7.766 [7.16, 8.50] years before the most recent sample (2006.175), corresponding to a root date of 1998.41, close to the 980-taxon reference (7.62 [7.21, 8.11]). The published 980-taxon analysis is used as an external timescale reference on a larger alignment, not a same-data model-comparison control. For the model-averaged 245-taxon and 980-taxon estimates, markers are posterior medians and bars are 95% highest-posterior-density intervals; the three family-conditional rates are shown as responsibility-weighted posterior means without intervals, since weighted credible intervals for model-conditional quantities are not reported (Supplementary Note 2). Terminal posterior model probabilities for this subset were *P*_S_ = 0.256, *P*_UC_ = 0.128, and *P*_AC_ = 0.615 (reported in the main text).

The structured-coalescent population-size summaries were broadly consistent with the original benchmark. The model-averaged posterior medians were 1.19 for Hong Kong (95% HPD 0.74– 1.78), 0.54 for New Zealand (95% HPD 0.34–0.78), and 1.54 for New York (95% HPD 1.01–2.17), whereas the full-data strict-clock analysis reported 1.67, 0.864, and 2.66, respectively. New Zealand was the smallest deme and New York the largest in both analyses, but the 245-taxon estimates were smaller than the full-data estimates for every deme. Because total tree length and total migration counts scale strongly with the number of sampled taxa, we do not compare those quantities directly between the 245- and 980-taxon analyses.

The H3N2 analysis shows that the hierarchical mixture can be applied in a demanding structured-coalescent setting, identifying the autocorrelated clock as the dominant relaxed-clock family while the strict clock retains non-negligible posterior probability, with an epidemic timescale that remains close to the published full-data strict-clock reference.

## 4 Discussion

The hierarchical clock mixture returns posterior support for the strict, UCLN, and autocorrelated clocks within a single Bayesian analysis, separating the strict-versus-relaxed choice from the within-relaxed choice and carrying both uncertainties into shared temporal summaries. Standard model-comparison approaches report support for competing clocks in a separate step from the downstream timescale summaries. The hierarchical mixture returns both within one posterior. When the data are only moderately informative, this avoids committing to a single clock prematurely; when they are strongly informative, the posterior concentrates on one clock family.

At the level of likelihood construction, the approach is analogous in spirit to mixture models that are already familiar in phylogenetics. Discrete-gamma site-rate models marginalise over latent rate categories at each site (Yang, 1994), and profile-mixture models such as CAT and finite C10– C60 mixtures marginalise over latent site profiles (Lartillot and Philippe, 2004; Si Quang et al., 2008). Those mixtures average over per-site latent categories within a fixed substitution and clock specification. The present construction instead averages over clock-family alternatives at the whole-tree likelihood level: the strict component does not depend on the branch-rate vector, whereas the relaxed component does. This distinction is what allows the top-level mixture to cross the strict– relaxed boundary without requiring a relaxed-clock prior to put positive mass on exactly equal branch rates.

Li and Drummond (2012) showed that Bayesian model averaging can be used within the relaxed-clock class to average over alternative branch-rate distributions rather than conditioning on a single relaxed-clock specification. The deterministic *u*-bridge described in Section 2.6 extends their quantile-parameterisation idea to cross-model moves between UCLN and the autocorrelated clock, with the Jacobian required when the MCMC state is the branch-rate vector itself. Marginal-likelihood estimators based on path sampling, stepping-stone sampling, and nested sampling remain the standard external route to clock-model comparison, and they have substantially improved over older estimators, but they still require separate analyses and a second inferential step in which biological interpretation is conditioned on one selected clock (Baele et al., 2012; Russel et al., 2019). The mixed clock of Lartillot et al. (2016) addresses a different question by blending uncorrelated and autocorrelated behaviour within a single generative model, whereas the present framework asks how posterior support should be allocated across separately interpretable clock families. Darlim and Höhna (2024) likewise show how analytical mixtures can compare competing priors within one MCMC analysis, but that construction does not by itself cross the strict-clock boundary because equal branch rates have zero probability under standard relaxed-clock priors. More recently, Panchaksaram et al. (2025) examined direct Bayesian selection between independent and autocorrelated relaxed clocks, including the effect of calibration misspecification on that comparison. Parallel developments within RevBayes pursue related model-comparison problems in a flexible graphical-model setting (Höhna et al., 2016), and recent proposal machinery in BEAST 2 has made shared-rate parameterisations increasingly practical (Douglas et al., 2021).

Under moderately informative simulation conditions, the posterior often did not collapse onto a single clock family even though the generating family was reliably retained in the credible set. This pattern reflects the information in the data: when data are informative enough to recover shared timescale parameters but not to distinguish decisively among nearby rate processes, carrying clock uncertainty forward is the appropriate posterior representation. The DENV-4 benchmark addressed a different question: whether a single-run posterior allocation would remain concordant with repeated evidence estimation. The mixture agreed with higher-effort nested sampling on the qualitative ranking of clock families while avoiding the run-to-run variability of independent marginal-likelihood estimates.

The four empirical benchmarks show that the mixture recovers the clock-family signal historically associated with each data set rather than being intrinsically biased toward a single family. The *rbcL* benchmark also showed that uncertainty outside the clock layer can propagate directly into clock-family support. Under a fixed JTT model, *rbcL* concentrated almost all posterior mass on the autocorrelated clock, consistent with its historical role in motivating autocorrelated-rate models (Thorne et al., 1998). Under an OBAMA site-model sensitivity analysis (Bouckaert, 2020), the strict clock remained effectively excluded, whereas both relaxed-clock families retained substantial posterior support: *P*_UC_ ≈ 0.409 and *P*_AC_ ≈ 0.591, with a modest preference for the autocorrelated model. This suggests that the richer site-model specification absorbed part of the heterogeneity that the fixed JTT analysis had to attribute to lineage-specific rate structure. The branch-level comparison further showed that UCLN-conditioned and autocorrelated-clock-conditioned mean rate multipliers were broadly concordant across retained branches (Fig. 3), indicating that the site-model sensitivity primarily altered posterior allocation between relaxed-clock families rather than producing strongly divergent branch-wise rate summaries. In the RSV-A G-gene benchmark, posterior mass concentrated almost entirely on UCLN, while the inferred viral timescale remained close to the estimates reported by Zlateva et al. (2004). The CCD0-MAP summary tree provided a branch-level view of the rate heterogeneity implied by this allocation (Fig. 4), making explicit that the earlier evidence against a single-rate molecular clock corresponds here to posterior support for UCLN rather than to diffuse uncertainty across relaxed-clock families. On the 245-taxon H3N2 subset, the relaxed class received most posterior mass with support concentrated on the autocorrelated family, and the strict clock retained non-negligible probability; because the subset changes both the data and the clock treatment relative to the published 980-taxon strict-clock analysis of Vaughan et al. (2014), we treat the timescale agreement as a benchmark check rather than a controlled estimate of the effect of clock choice. These examples illustrate that clock-model support and downstream timescale sensitivity are related but not identical (Ho and Duchêne, 2014; Duchêne et al., 2014): a preferred clock model may change while some shared summaries remain robust, and robustness of a particular summary is not itself a reason to fix the clock in advance.

The current implementation considers three clock families: one strict clock and two relaxed clocks. Many other biologically plausible rate processes are not included. It is also not a mixed-clock generative model in which different regions of the tree can follow different rate processes. The empirical data sets were chosen as benchmarks with distinct inferential roles, not as a broad survey of rate variation across the Tree of Life. The simulation study likewise covers only moderately informative data. In those simulations, increasing the amount of data through more sites, more leaves, or higher but non-saturated mutation rates concentrated posterior probability on the generating clock family. Averaging over clock families does not make the analysis robust to calibration misspecification: severe misspecification can distort posterior allocation within the relaxed-clock class itself (Panchaksaram et al., 2025).

Several directions could extend this framework. One is to enlarge the clock-model space to include additional rate processes, such as mixed or additive relaxed clocks (Lartillot et al., 2016; Didelot et al., 2021). This extension would require generalising both cross-model operators. The Gibbs update adapts naturally to a *K*-way categorical draw, whereas the current UC–AC bridge relies on both relaxed-clock priors being represented through Gaussian latent variables. Extending the bridge to non-Gaussian branch-rate priors would likely require an explicit uniform-quantile parameterisation (Li and Drummond, 2012). A second direction is to combine clock-model uncertainty with uncertainty in demographic model classes, such as the parametric-coalescent model averaging of Xu et al. (2025). Because the clock prior and the tree prior enter the posterior as separate factors, the two averaging schemes are modular and can in principle already be composed within a single BEAST 2 analysis simply by selecting both in BEAUti, so that clock-model and tree-prior uncertainty are propagated jointly rather than one being fixed while the other is averaged over. What remains is therefore less a question of feasibility than of confirming that the combined operator scheme mixes adequately and that the joint model-averaged posterior behaves as intended, and of demonstrating the practical benefit on real data. Finally, relative comparison among candidate clock models is not the same as absolute model adequacy or temporal-signal adequacy. Those questions can be assessed independently by posterior-predictive checks (Duchêne et al., 2015) and BETS (Duchene et al., 2020). The main conclusion is simpler: clock-model uncertainty in Bayesian phylogenetic dating need not be resolved through repeated evidence estimation across separate runs; it can be represented directly within the posterior and propagated into the divergence-time statements that are finally reported.

## Acknowledgements

We thank the Centre for Computational Evolution at the University of Auckland for computational resources and helpful discussions.

## Data Availability

Analysis-ready empirical alignments, sampling-date metadata, BEAST 2 XML files, LinguaPhylo simulation scripts, nested-sampling configuration files, post-processing scripts, figure-generation scripts, posterior summary files, and processed data files needed to reproduce the analyses reported in this study are available to reviewers through the temporary Dryad Reviewer URL: http://datadryad.org/share/LINK_NOT_FOR_PUBLICATION/gBkczj_DgCazSqSaJhtLHaYO8FZtd066CPxQwDypFUI. The `RelaxClockAveraging` BEAST package source code, including the implementation of the hierarchical clock-mixture model and associated example/template files, is archived in Zenodo: https://doi.org/10.5281/zenodo.20684860.

Development versions of the software and analysis repository are available at https://github.com/yxu927/RelaxClockAveraging and https://github.com/yxu927/mixture_data. Previously published sequence data analysed in this study are available from the original sources cited in the manuscript; the analysis-ready versions used here are included in the Dryad archive.

