## Supplemental file for "A hierarchical clock-mixture model for Bayesian phylogenetic dating"

### 1 Supplementary Note 1. Full prior specification within the relaxed superclass

#### 1.1 Within-relaxed indicator and full joint posterior

Within the relaxed superclass, uncertainty between the uncorrelated and autocorrelated rate processes is represented by a binary indicator $z \in \{0, 1\}$, where $z = 0$ selects the uncorrelated model and $z = 1$ selects the autocorrelated model. Conditional on entering the relaxed superclass, the two submodels receive equal prior mass,

$$\Pr(z = 0) = \Pr(z = 1) = \frac{1}{2}. \tag{1}$$

Let $\sigma_{\mathrm{U}} > 0$ denote the log-scale spread parameter under the uncorrelated model, $\nu_{\mathrm{A}} > 0$ the diffusion variance under the autocorrelated model, and $x_0 \in \mathbb{R}$ an optional root log-rate anchor. The full posterior is

$$\begin{aligned} p(T, \mu, \boldsymbol{r}, \phi, \psi, z, \sigma_{\mathrm{U}}, \nu_{\mathrm{A}}, x_0 \mid D) \propto\ & L_{\mathrm{top}}(D \mid T, \mu, \boldsymbol{r}, \phi) \\ & \times \pi(\boldsymbol{r} \mid T, z, \sigma_{\mathrm{U}}, \nu_{\mathrm{A}}, x_0)\, \pi(z) \\ & \times \pi(T \mid \psi)\, \pi(\psi)\, \pi(\mu)\, \pi(\phi)\, \pi(\sigma_{\mathrm{U}})\, \pi(\nu_{\mathrm{A}})\, \pi(x_0). \end{aligned} \tag{2}$$

When $x_0$ is fixed rather than estimated, $\pi(x_0)$ is understood as a degenerate prior and $x_0$ is omitted from posterior summaries.

The top-level mixture likelihood is

$$L_{\mathrm{top}}(D \mid T, \mu, \boldsymbol{r}, \phi) = w_{\mathrm{S}} L_{\mathrm{S}}(D \mid T, \mu, \phi) + w_{\mathrm{R}} L_{\mathrm{R}}(D \mid T, \mu, \boldsymbol{r}, \phi).$$

The branch-rate vector $\boldsymbol{r}$, the within-relaxed indicator $z$, and the relaxed-clock hyperparameters are present in the augmented state at all MCMC iterations. When the strict likelihood component contributes to the top-level mixture, these variables are auxiliary variables with proper priors. Integrating over them gives one for the strict component, so the strict-clock marginal contribution is $w_{\mathrm{S}}Z_{\mathrm{S}}$, whereas the relaxed-clock marginal contribution is

$$w_{\mathrm{R}}\left(\eta_{\mathrm{UC}}Z_{\mathrm{UC}}+\eta_{\mathrm{AC}}Z_{\mathrm{AC}}\right).$$

Thus the terminal prior model masses are

$$\Pr(M_{\mathrm{S}})=w_{\mathrm{S}},\qquad \Pr(M_{\mathrm{UC}})=w_{\mathrm{R}}\eta_{\mathrm{UC}},\qquad \Pr(M_{\mathrm{AC}})=w_{\mathrm{R}}\eta_{\mathrm{AC}}.$$

With $w_{\mathrm{S}}=1/3$, $w_{\mathrm{R}}=2/3$, and $\eta_{\mathrm{UC}}=\eta_{\mathrm{AC}}=1/2$, the three terminal clock families have equal prior mass.

##### 1.2 Uncorrelated lognormal prior

Under the uncorrelated lognormal clock (Drummond et al., 2006),

$$r_b \overset{\mathrm{iid}}{\sim} \mathrm{LogNormal}\left(-\tfrac{1}{2}\sigma_{\mathrm{U}}^2,\sigma_{\mathrm{U}}^2\right),\qquad b\in B, \tag{3}$$

so that $\mathbb{E}[r_b]=1$. Evaluated on the positive rate scale, the joint density is

$$f_{\mathrm{UC}}(\boldsymbol{r}\mid\sigma_{\mathrm{U}})=\prod_{b\in B}\frac{1}{r_b\,\sigma_{\mathrm{U}}\sqrt{2\pi}}\exp\left[-\frac{\left(\log r_b+\frac{1}{2}\sigma_{\mathrm{U}}^2\right)^2}{2\sigma_{\mathrm{U}}^2}\right]. \tag{4}$$

The factor $1/r_b$ is the Jacobian of the log transformation.

##### 1.3 Autocorrelated lognormal prior

Under the autocorrelated lognormal clock (Thorne et al., 1998; Kishino et al., 2001), rates evolve as a rooted first-order Markov process on the log scale. For each non-root branch $b$ with parent branch $\mathrm{pa}(b)$ and duration $t_b>0$,

$$\log r_b\mid\log r_{\mathrm{pa}(b)}\sim\mathcal{N}\left(\log r_{\mathrm{pa}(b)}-\tfrac{1}{2}\nu_{\mathrm{A}}t_b,\ \nu_{\mathrm{A}}t_b\right). \tag{5}$$

For branches immediately descending from the root, $\log r_{\mathrm{pa}(b)}$ is replaced by $x_0$. The mean correction ensures

$$\mathbb{E}[r_b\mid r_{\mathrm{pa}(b)}]=r_{\mathrm{pa}(b)}, \tag{6}$$

so that the process is centred on multiplicative inheritance rather than upward drift on the rate scale.

The corresponding rate-space density is

$$f_{\mathrm{AC}}(\boldsymbol{r}\mid T,\nu_{\mathrm{A}},x_0)=\prod_{b\in B}\frac{1}{r_b\sqrt{2\pi\nu_{\mathrm{A}}t_b}}\exp\left[-\frac{\left(\log r_b-m_b\right)^2}{2\nu_{\mathrm{A}}t_b}\right], \tag{7}$$

where

$$m_b = \begin{cases} x_0 - \frac{1}{2}\nu_{\mathrm{A}} t_b, & \text{if } \mathrm{pa}(b) \text{ is the root}, \\ \log r_{\mathrm{pa}(b)} - \frac{1}{2}\nu_{\mathrm{A}} t_b, & \text{otherwise.} \end{cases} \tag{8}$$

Again, the factor $1/r_b$ is the Jacobian of the log transformation.

##### 1.4 Implementation safeguard for short branches

In the current implementation, the autocorrelated prior assigns $-\infty$ log density whenever any non-root branch satisfies $t_b \leq t_{\min}$, where $t_{\min} > 0$ is a small numerical threshold (default $10^{-12}$ in user-specified time units). This is a numerical safeguard against degenerate diffusion variances on zero-length or nearly zero-length branches rather than a defining feature of the idealised autocorrelated model.

##### 1.5 Variable-selection prior on the shared rate vector

The prior on the shared rate vector is therefore

$$\pi(\boldsymbol{r} \mid T, z, \sigma_{\mathrm{U}}, \nu_{\mathrm{A}}, x_0) = \begin{cases} f_{\mathrm{UC}}(\boldsymbol{r} \mid \sigma_{\mathrm{U}}), & z = 0, \\ f_{\mathrm{AC}}(\boldsymbol{r} \mid T, \nu_{\mathrm{A}}, x_0), & z = 1. \end{cases} \tag{9}$$

The indicator changes only the prior on the (shared) rate vector $\boldsymbol{r}$ and therefore does not induce a dimension change in the state space.

#### 2 Supplementary Note 2. Posterior model probabilities, Bayes factors, and conditional reporting

##### 2.1 Iteration-wise responsibilities and posterior support

Because the top-level strict-versus-relaxed indicator is marginalised analytically, posterior support for the strict and relaxed superclasses is recovered from the iteration-wise conditional probabilities of each clock family. We refer to these per-state conditional probabilities as responsibilities (sensu Bishop and Nasrabadi, 2006): at each MCMC state, the responsibility of a clock family is the posterior probability that this family is the active one, given the current continuous parameters. At retained iteration $m$, let $L_{\mathrm{S}}^{(m)}$ and $L_{\mathrm{R}}^{(m)}$ denote the strict- and relaxed-clock likelihoods evaluated at the current state. The top-level responsibilities are

$$r_{\mathrm{R}}^{(m)} = \frac{w_{\mathrm{R}} L_{\mathrm{R}}^{(m)}}{w_{\mathrm{S}} L_{\mathrm{S}}^{(m)} + w_{\mathrm{R}} L_{\mathrm{R}}^{(m)}}, \qquad r_{\mathrm{S}}^{(m)} = 1 - r_{\mathrm{R}}^{(m)}.$$

Within the relaxed superclass, the UCLN and autocorrelated priors are compared on the same branch-rate vector. Let $f_{\mathrm{UC}}(\boldsymbol{r}^{(m)} \mid \sigma_{\mathrm{U}}^{(m)})$ and $f_{\mathrm{AC}}(\boldsymbol{r}^{(m)} \mid T^{(m)}, \nu_{\mathrm{A}}^{(m)}, x_0^{(m)})$ denote the two relaxed-clock prior densities. The corresponding within-relaxed responsibilities are

$$\tau_{\mathrm{UC}}^{(m)} = \frac{\eta_{\mathrm{UC}} f_{\mathrm{UC}}(\boldsymbol{r}^{(m)} \mid \sigma_{\mathrm{U}}^{(m)})}{\eta_{\mathrm{UC}} f_{\mathrm{UC}}(\boldsymbol{r}^{(m)} \mid \sigma_{\mathrm{U}}^{(m)}) + \eta_{\mathrm{AC}} f_{\mathrm{AC}}(\boldsymbol{r}^{(m)} \mid T^{(m)}, \nu_{\mathrm{A}}^{(m)}, x_0^{(m)})}, \qquad \tau_{\mathrm{AC}}^{(m)} = 1 - \tau_{\mathrm{UC}}^{(m)}.$$

The Rao–Blackwellized per-iteration terminal responsibilities are therefore

$$p_{\mathrm{S}}^{(m)} = r_{\mathrm{S}}^{(m)}, \qquad p_{\mathrm{UC}}^{(m)} = r_{\mathrm{R}}^{(m)} \tau_{\mathrm{UC}}^{(m)}, \qquad p_{\mathrm{AC}}^{(m)} = r_{\mathrm{R}}^{(m)} \tau_{\mathrm{AC}}^{(m)}.$$

By construction,

$$p_{\mathrm{S}}^{(m)} + p_{\mathrm{UC}}^{(m)} + p_{\mathrm{AC}}^{(m)} = 1.$$

Posterior model probabilities are estimated by Monte Carlo averaging:

$$\widehat{\Pr}(M_k \mid D) = \frac{1}{N_{\mathrm{iter}}} \sum_{m=1}^{N_{\mathrm{iter}}} p_k^{(m)}, \qquad k \in \{\mathrm{S}, \mathrm{UC}, \mathrm{AC}\}.$$

A sampled-indicator estimator, $p_{\mathrm{UC}}^{(m)} = r_{\mathrm{R}}^{(m)} \mathbf{1}\{z^{(m)} = 0\}$ and $p_{\mathrm{AC}}^{(m)} = r_{\mathrm{R}}^{(m)} \mathbf{1}\{z^{(m)} = 1\}$, is also unbiased when $z$ is sampled from its full conditional, but the Rao–Blackwellized estimator above uses the full conditional information at each retained state and has lower Monte Carlo variance.

#### 2.2 Bayes factors under equal and unequal prior masses

Under the family-balanced prior calibration,

$$\Pr(M_{\mathrm{S}}) = \Pr(M_{\mathrm{UC}}) = \Pr(M_{\mathrm{AC}}) = \frac{1}{3}. \tag{10}$$

Pairwise Bayes factors therefore reduce to posterior-odds ratios,

$$BF_{ij} = \frac{\widehat{\Pr}(M_i \mid D)}{\widehat{\Pr}(M_j \mid D)}, \qquad i \neq j. \tag{11}$$

If alternative prior weights are used at either level of the hierarchy, the correct prior-odds adjustment is

$$BF_{ij} = \frac{\widehat{\Pr}(M_i \mid D)/\widehat{\Pr}(M_j \mid D)}{\Pr(M_i)/\Pr(M_j)}. \tag{12}$$

For numerical stability, especially when posterior support for one family is close to zero, Bayes factors should be evaluated on the log scale.

##### 2.3 Conditional summaries of submodel-specific parameters

Parameters that belong to only one relaxed-clock submodel should be summarised conditionally on that submodel. Let $\theta_{\mathrm{UC}}$ denote a UCLN-specific parameter, such as $\sigma_{\mathrm{U}}$, and let $\theta_{\mathrm{AC}}$ denote an autocorrelated-clock-specific parameter, such as $\nu_{\mathrm{A}}$ or, when estimated, $x_0$. Using the terminal responsibilities $p_k^{(m)}$ defined above, the conditional posterior mean under each submodel is estimated as

$$\widehat{\mathbb{E}}(\theta_{\mathrm{UC}} \mid D, M_{\mathrm{UC}}) = \frac{\sum_{m=1}^{N_{\mathrm{iter}}} p_{\mathrm{UC}}^{(m)} \theta_{\mathrm{UC}}^{(m)}}{\sum_{m=1}^{N_{\mathrm{iter}}} p_{\mathrm{UC}}^{(m)}}, \qquad \widehat{\mathbb{E}}(\theta_{\mathrm{AC}} \mid D, M_{\mathrm{AC}}) = \frac{\sum_{m=1}^{N_{\mathrm{iter}}} p_{\mathrm{AC}}^{(m)} \theta_{\mathrm{AC}}^{(m)}}{\sum_{m=1}^{N_{\mathrm{iter}}} p_{\mathrm{AC}}^{(m)}}.$$

Weighting by $p_{\mathrm{UC}}^{(m)}$ (respectively $p_{\mathrm{AC}}^{(m)}$) downweights states in which that family is inactive—where the corresponding parameter is effectively a draw from its prior—so the conditional mean is informed almost entirely by states in which the family is active.

We report only this responsibility-weighted conditional mean for submodel-specific parameters. We do not summarise them by weighted medians or weighted credible intervals: because the responsibilities are unequal across states, the weighted sample has a reduced effective sample size, and weighted quantiles are less well justified than the weighted mean. When one relaxed-clock family carries almost all posterior support—for example UCLN in the RSV-A analysis, where $P_{\mathrm{UC}} \approx 0.99$—the responsibilities are effectively uniform and the conditional summary reduces to an ordinary posterior summary restricted to that family.

#### 3 Supplementary Note 3. Simulation protocol

##### 3.1 Simulation design

Simulation-based validation was performed under correct model specification using a stratified design. The purpose of the simulation study was to validate implementation correctness and posterior calibration under the model class used for inference, rather than to assess robustness to model misspecification. We generated 100 replicate alignments under each of the three terminal clock families: strict, UCLN, and autocorrelated. Thus the terminal clock family was fixed within each simulation stratum, while the continuous parameters were drawn from the same conditional priors used in the corresponding component of the inference model.

Across all three strata, 30 heterochronously sampled taxa were placed at one-third-unit intervals. Genealogies were drawn from a constant-size coalescent with

$$\Theta \sim \mathrm{LogNormal}(1.2, 0.6),$$

and alignments of 1500 nucleotides were simulated under JC69 with four-category discrete-gamma site-rate variation,

$$\gamma \sim \mathrm{LogNormal}(-1.0, 0.7).$$

The global mean rate was drawn as

$$\mu \sim \text{LogNormal}(-3.4, 0.5).$$

For strict-clock simulations, all branches shared the global mean rate. For UCLN simulations, branch-specific relative rates were drawn under the UCLN prior with

$$\sigma_{\text{U}} \sim \text{LogNormal}(-1.1, 0.6).$$

For autocorrelated-clock simulations, log rates evolved along the tree with diffusion variance

$$\nu_{\text{A}} \sim \text{LogNormal}(-4, 1.0),$$

and the root log-rate anchor was fixed at $x_0 = 0$. Each simulated alignment was then analysed under the full hierarchical clock-mixture model with the same terminal prior masses and inference priors as in the empirical analyses.

#### 3.2 LPhy simulation script

Each replicate dataset was generated using LPhyStudio within the LinguaPhylo framework (Drummond et al., 2023). Listing S1 gives the common prior-predictive template for the hierarchical clock model. For the stratified simulation study reported in the main text, however, the terminal clock family was fixed within each simulation stratum rather than drawn from the top-level model prior. Specifically, strict-clock replicates used the strict likelihood component, UCLN replicates used the relaxed component with the within-relaxed indicator fixed to UCLN, and autocorrelated replicates used the relaxed component with the within-relaxed indicator fixed to the autocorrelated clock. All continuous parameters were drawn from the same conditional priors used in the corresponding component of the inference model.

Thus Listing S1 should be read as the common model template, not as a literal single prior-predictive script in which the terminal family was randomly drawn for the 300 stratified validation replicates.

Listing 1: Common LPhy template used for simulation-based validation. For the stratified simulations reported in the main text, the terminal clock indicators were fixed within each simulation stratum: strict for the strict-clock stratum, relaxed with the UCLN indicator for the UCLN stratum, and relaxed with the autocorrelated indicator for the autocorrelated stratum. Continuous parameters were drawn from the same conditional priors shown in the template.

```
data {
    dt   = 1.0/3.0;
    tmax = 10.0;
    ages = arange(start=0.0, stop=tmax, step=dt);
    n    = length(ages);
    names = rangeInt(start=1, end=n);
    taxa  = taxa(names=names, ages=ages);
```

```
    nchar = 1500;
}

model {
    // Coalescent tree prior
    Theta ~ LogNormal(meanlog=1.2, sdlog=0.6);
    psi   ~ Coalescent(taxa=taxa, theta=Theta);

    // Site-rate variation (JC69 + discrete gamma)
    gamma ~ LogNormal(meanlog=-1.0, sdlog=0.7);
    siteRates ~ DiscretizeGamma(shape=gamma, ncat=4, replicates=nchar);

    // Global mean rate
    clockRate ~ LogNormal(meanlog=-3.4, sdlog=0.5);

    // Relaxed-clock hyperparameters
    sigmaUCLN  ~ LogNormal(meanlog=-1.1, sdlog=0.6);
    rootLogRate = 0.0;
    sigma2     ~ LogNormal(meanlog=-4, sdlog=1.0);

    // Level-2 draw: UCLN vs. autocorrelated relaxed clock (equal weights)
    w_relax = [1.0/2, 1.0/2];
    i_relax ~ Categorical(p=w_relax);
    rawRates ~ SVSRawBranchRates(tree=psi, indicator=i_relax,
                                 ucldStdev=sigmaUCLN, sigma2=sigma2,
                                 rootLogRate=rootLogRate);
    branchRates = SharedRatesClock(tree=psi, rates=rawRates,
                                  meanRate=clockRate, normalize=true);

    // Level-1 draw: strict (1/3) vs. relaxed (2/3)
    w_top = [1.0/3, 2.0/3];
    i_top ~ Categorical(p=w_top);
    D_strict  ~ PhyloCTMC(L=nchar, Q=jukesCantor(),
                          siteRates=siteRates, mu=clockRate, tree=psi);
    D_relaxed ~ PhyloCTMC(L=nchar, Q=jukesCantor(),
                          siteRates=siteRates, tree=psi,
                          branchRates=branchRates);
    D ~ MixturePhyloCTMC(comp1=D_strict, comp2=D_relaxed,
                         index=i_top, weights=w_top);

    // Deterministic summaries
    length = psi.treeLength();
    height = psi.rootAge();
}
```

The two-level indicator structure (`i_top` and `i_relax`) reflects the hierarchical mixture described in the main text: $i_{\text{top}} \in \{0, 1\}$ selects between the strict and relaxed families, and the subordinate draw $i_{\text{relax}} \in \{0, 1\}$ is relevant only when the relaxed family is active. Branch rates are assembled by `SVSRawBranchRates`, which routes to either the uncorrelated lognormal or the autocorrelated lognormal process according to `i_relax`, and then rescaled to the specified global mean rate by `SharedRatesClock`. Sequence data are generated from the mixture distribution `MixturePhyloCTMC`, which selects the active likelihood component via `i_top`. The deterministic nodes `length` and `height` record the true tree length and root age for use in subsequent coverage assessment.

##### 3.3 Additional simulation condition (shallower, lower-information)

Listing 2: Under a lower substitution rate ($\mu \sim \text{LogNormal}(-5, 0.5)$) and shorter alignments (600 nucleotides), placing the data in a lower-information regime (median $\mu H = 0.10$ substitutions site$^{-1}$, central 95% range 0.04–0.29).

```
data {
    dt   = 1.0/3.0;
    tmax = 10.0;
    ages = arange(start=0.0, stop=tmax, step=dt);
    n    = length(ages);
    names = rangeInt(start=1, end=n);
    taxa  = taxa(names=names, ages=ages);
    nchar = 600;
}

model {
    // Coalescent tree prior
    Theta ~ LogNormal(meanlog=1.2, sdlog=0.6);
    psi   ~ Coalescent(taxa=taxa, theta=Theta);

    // Site-rate variation (JC69 + discrete gamma)
    gamma ~ LogNormal(meanlog=-1.0, sdlog=0.7);
    siteRates ~ DiscretizeGamma(shape=gamma, ncat=4, replicates=nchar);

    // Global mean rate
    clockRate ~ LogNormal(meanlog=-5.0, sdlog=0.5);

    // Relaxed-clock hyperparameters
    sigmaUCLN  ~ LogNormal(meanlog=-1.1, sdlog=0.6);
    rootLogRate = 0.0;
    sigma2     ~ LogNormal(meanlog=-4, sdlog=1.0);

    // Level-2 draw: UCLN vs. autocorrelated relaxed clock (equal weights)
    w_relax = [1.0/2, 1.0/2];
    i_relax ~ Categorical(p=w_relax);
    rawRates ~ SVSRawBranchRates(tree=psi, indicator=i_relax,
                                 ucldStdev=sigmaUCLN, sigma2=sigma2,
                                 rootLogRate=rootLogRate);
```

```
    branchRates = SharedRatesClock(tree=psi, rates=rawRates,
                                   meanRate=clockRate, normalize=true);

    // Level-1 draw: strict (1/3) vs. relaxed (2/3)
    w_top = [1.0/3, 2.0/3];
    i_top ~ Categorical(p=w_top);
    D_strict  ~ PhyloCTMC(L=nchar, Q=jukesCantor(),
                          siteRates=siteRates, mu=clockRate, tree=psi);
    D_relaxed ~ PhyloCTMC(L=nchar, Q=jukesCantor(),
                          siteRates=siteRates, tree=psi,
                          branchRates=branchRates);
    D ~ MixturePhyloCTMC(comp1=D_strict, comp2=D_relaxed,
                         index=i_top, weights=w_top);

    // Deterministic summaries
    length = psi.treeLength();
    height = psi.rootAge();
}
```

##### 3.4 Results for the additional simulation condition

In this condition we lowered the substitution rate to $\mu \sim \text{LogNormal}(-5, 0.5)$ and shortened alignments to 600 nucleotides, lowering the median mean-rate-scaled root age from $\mu H = 0.50$ in the main study (Figure 2 in the main text) to $\mu H = 0.10$. Reducing the information content of the data in this way left coverage of the true clock family unchanged but markedly increased the residual model-level uncertainty. The generating clock family was again retained in the 95% posterior model set in all 300 replicates (Fig. S1A; Table S1). The credible-set-size distribution, however, shifted from predominantly singleton sets in the higher-information main study (213 of 300 replicates, 71%) to predominantly two-model sets here (234 of 300 replicates, 78%). The shift was most pronounced for strict-clock simulations, which yielded a singleton set in 36% of replicates in the main study but in none of the 100 replicates under the lower-information regime, reflecting that the strict-versus-relaxed distinction is the hardest to resolve when the data are least informative. Model-averaged estimation of the shared timescale parameters nevertheless remained accurate, with 95% HPD coverage between 91% and 100% for the overall rate, root age, and tree length (Table S1).

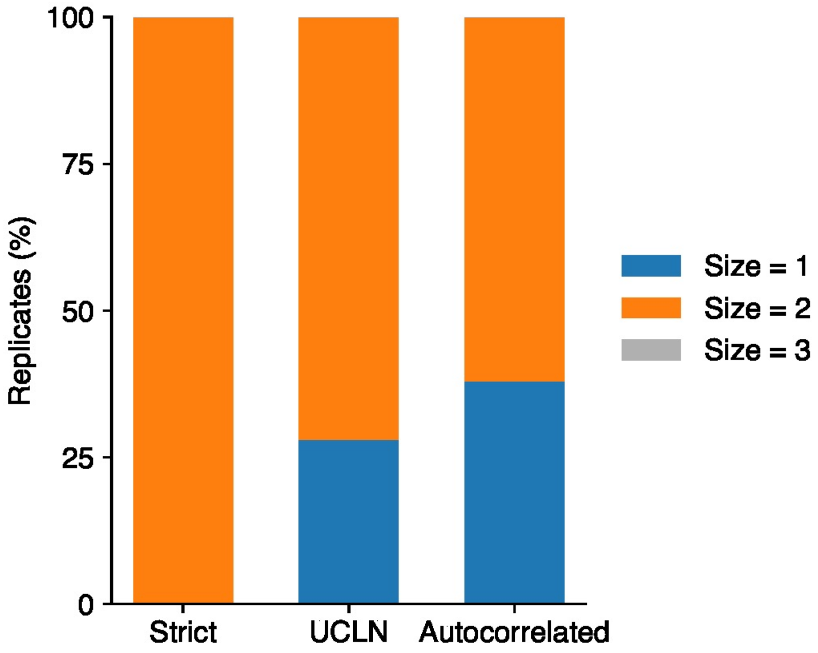

Figure S1: Stratified simulation study of clock-model recovery under the lower-information regime ($\mu \sim \text{LogNormal}(-5, 0.5)$, 600-nucleotide alignments, median $\mu H = 0.10$), across 300 replicates. This is the low-information counterpart to the higher-information main study of Figure 2 in the main text (median $\mu H = 0.50$, 1500-nucleotide alignments). **(A)** Empirical coverage of the 95% posterior model credible set for each generating clock model. The dashed line indicates the nominal 95% level. **(B)** Distribution of credible set sizes. Most replicates retained two clock models and none retained all three—in contrast to the higher-information main study, where most replicates yielded a singleton set—reflecting conservative rather than sharply calibrated behaviour in this lower-information regime.

Table S1: Summary of simulation-based validation under the lower-information simulation condition.

| | | Credible set size (%) | | | HPD coverage (%) | | |
|---|---|---|---|---|---|---|---|
| True model | In 95% set (%) | Size 1 | Size 2 | Size 3 | Clock rate | Root age | Tree length |
| Strict | 100 | 0 | 100 | 0 | 100 | 94 | 92 |
| UCLN | 100 | 28 | 72 | 0 | 96 | 93 | 96 |
| Autocorrelated | 100 | 38 | 62 | 0 | 91 | 93 | 95 |

### 4 Supplementary Note 4. MCMC operator derivations and diagnostics

#### 4.1 Cross-model proposals for the within-relaxed indicator

Sampling of the binary indicator $z \in \{0, 1\}$ results in a non-trivial mixing problem in the within-relaxed part of the model. A naive flip of $z$ without adjusting the branch-rate vector $\boldsymbol{r}$ or the hyperparameters $(\sigma_{\mathrm{U}}, \nu_{\mathrm{A}})$ typically proposes a state whose density under the new prior is vanishingly small, because a rate configuration drawn from one prior process rarely looks natural under the other. Ordinary Metropolis–Hastings flips of $z$ therefore accept very rarely, and the chain becomes stuck on whichever clock family it started in. We used two specialised cross-model operators for the within-relaxed indicator: an exact Gibbs update of $z$, and a deterministic bridge proposal that flips $z$ while coherently remapping the shared branch-rate vector between the UCLN and autocorrelated parameterisations. Table S2 summarises which state variables each operator proposes to change.

**Strategy 1: Gibbs update on the indicator.** The operator `IndicatorGibbsOperator` proposes $z$ from its exact full conditional given the current rates and hyperparameters,

$$\Pr(z = k \mid \boldsymbol{r}, T, \sigma_{\mathrm{U}}, \nu_{\mathrm{A}}, x_0) \;\propto\; \eta_k\, f_k(\boldsymbol{r} \mid T, \sigma_{\mathrm{U}}, \nu_{\mathrm{A}}, x_0), \qquad k \in \{0, 1\}, \tag{13}$$

where $f_0 = f_{\mathrm{UC}}$ and $f_1 = f_{\mathrm{AC}}$ are the UCLN and autocorrelated priors on the branch-rate vector and $\eta_k$ are the prior weights on the two relaxed-clock models. The phylogenetic likelihood does not depend on $z$ directly (both models act on the same $\boldsymbol{r}$), so the Hastings ratio cancels and the move is accepted with probability one. This update is exact conditional on $\boldsymbol{r}$ and the hyperparameters, but it does not itself reconfigure $\boldsymbol{r}$ or the hyperparameters to be plausible under the proposed model, which motivates the complementary rate-bridge proposal below. The indicator-Gibbs construction is standard in Bayesian variable and model selection (George and McCulloch, 1993; Kuo and Mallick, 1998). For the Gibbs update we compute the unnormalised log weights

$$a_0 = \log \eta_{\mathrm{UC}} + \log f_{\mathrm{UC}}(\boldsymbol{r} \mid \sigma_{\mathrm{U}}), \qquad a_1 = \log \eta_{\mathrm{AC}} + \log f_{\mathrm{AC}}(\boldsymbol{r} \mid T, \nu_{\mathrm{A}}, x_0),$$

where $\eta_{\mathrm{UC}}$ and $\eta_{\mathrm{AC}}$ are the prior weights on the two relaxed-clock models. Note that $f_{\mathrm{UC}}$ does not condition on the tree $T$ because UCLN treats branch rates as independent and identically distributed draws from a lognormal distribution, so the prior density on $\boldsymbol{r}$ depends only on the hyperparameter $\sigma_{\mathrm{U}}$; the autocorrelated prior, by contrast, conditions each branch rate on the rate of its parent node and the branch time, and therefore requires $T$ and the root log-rate $x_0$. The normalised full conditional probabilities are evaluated using a log-sum-exp calculation,

$$\tau_k = \Pr(z = k \mid \boldsymbol{r}, T, \sigma_{\mathrm{U}}, \nu_{\mathrm{A}}, x_0) = \exp\{a_k - \operatorname{logsumexp}(a_0, a_1)\}, \qquad k \in \{0, 1\}.$$

The proposal draws $z' \sim \text{Categorical}(\tau_0, \tau_1)$. Because the proposal distribution is the full conditional, the returned Hastings correction is

$$\log H = \log \tau_z - \log \tau_{z'},$$

which exactly cancels the target-density ratio involving $z$. The move is therefore accepted with probability one, conditional on the current branch-rate vector and hyperparameters. In all analyses reported here we used $\eta_{\text{UC}} = \eta_{\text{AC}} = 1/2$.

**Strategy 2: rate-changing proposals.** The second operator flips $z$ and also changes the branch-rate vector, so the phylogenetic likelihood changes across the switch.

`UCACSwitchBridgeOperator` replaces the fresh prior draw with a deterministic quantile-preserving remap. Both relaxed priors can be written as a non-centered parameterisation (Papaspiliopoulos et al., 2007) of the branch-rate vector in terms of a shared latent standard-normal vector $\mathbf{u}$: under UCLN, $\log r_b = -\sigma_{\text{U}}^2/2 + \sigma_{\text{U}} u_b$ with $u_b \sim \mathcal{N}(0,1)$; under the autocorrelated clock, $\log r_b = \log r_{\text{pa}(b)} - \frac{1}{2}\nu_{\text{A}} t_b + \sqrt{\nu_{\text{A}} t_b}\, u_b$, again with $u_b \sim \mathcal{N}(0,1)$. The operator reads $\mathbf{u}$ off the current $\boldsymbol{r}$ by inverting the current model's map, flips $z$, and pushes the same $\mathbf{u}$ forward through the new model's map to obtain $\boldsymbol{r}'$. This is the quantile-parameterisation idea of Li and Drummond (2012) applied to the UC/AC boundary: each branch retains the same Gaussian-quantile position under its new prior, while the map from quantile to rate changes with $z$. Because the state here is $\boldsymbol{r}$ rather than $\mathbf{u}$, the move is a change of variables that incurs a Jacobian. Writing $|B|$ for the number of non-root branches, the log-Jacobian is

$$\log |J_{\text{UC}\to\text{AC}}| \;=\; \sum_{b\in B} \log \frac{r'_b}{r_b} \;+\; \frac{1}{2}\sum_{b\in B} \log(\nu_{\text{A}} t_b) \;-\; |B| \log \sigma_{\text{U}}, \tag{14}$$

with the sign of the second and third terms reversed for the AC→UC direction, giving

$$\log |J_{\text{AC}\to\text{UC}}| \;=\; \sum_{b\in B} \log \frac{r'_b}{r_b} \;-\; \frac{1}{2}\sum_{b\in B} \log(\nu_{\text{A}} t_b) \;+\; |B| \log \sigma_{\text{U}}. \tag{15}$$

Because the proposal is fully deterministic, $\log |J|$ is the entire Hastings correction, and the log-acceptance ratio is $\log \tilde{p}(\text{new}) - \log \tilde{p}(\text{old}) + \log |J|$ where $\tilde{p}$ is the unnormalised posterior density. The strategy directly addresses prior mismatch on large trees: $\boldsymbol{r}'$ has the same whitened-residual structure as $\boldsymbol{r}$, so it tends to sit in a high-posterior region under the new prior as well.

Table S2: Cross-model MCMC operators for the within-relaxed indicator $z$. A tick indicates that the corresponding state variable is proposed to change. The final column gives the Hastings correction beyond the target-density ratio.

| Operator | $z$ | $\boldsymbol{r}$ | Type | Hastings term |
|---|---|---|---|---|
| `IndicatorGibbsOperator` | ✓ | — | Gibbs | $\log \tau_z - \log \tau_{z'}$; acceptance = 1 |
| `UCACSwitchBridgeOperator` | ✓ | ✓ | MH (deterministic) | Eqs. 14 & 15 |

In the analyses reported in the main text, the cross-model operators were combined with standard multiplicative scale proposals on individual rates and on subtree blocks of $\boldsymbol{r}$, and with a within-model subtree proposal tailored to the autocorrelated component that is described in Section 4.2.

In all empirical analyses using this two-operator configuration, the `IndicatorGibbsOperator` and `UCACSwitchBridgeOperator` were assigned weights of 15 and 10 in the XML, respectively, which determine how often the MCMC algorithm selects them to do a proposal. The deterministic UC–AC bridge showed more data-dependent behaviour (Table S3): it accepted frequently in the DENV-4 benchmark, had non-negligible acceptance in the H3N2 structured-coalescent analysis, and accepted rarely in the *rbcL* and RSV-A analyses. We therefore used the Gibbs update as the reliable routine update of the within-relaxed indicator, while retaining the bridge as a complementary long-range proposal that can move the branch-rate vector coherently between the UCLN and autocorrelated parameterisations.

The Gibbs update accepts every proposal by construction, so we instead report how often the indicator $z$ actually changes (Table S3); for the deterministic bridge an accepted move always flips $z$, so its acceptance fraction is already its switch rate.

Table S3: Empirical mixing of the within-relaxed indicator $z$. The second column is the realised probability that $z$ differs between successive logged states (every $10^3$ states, 10% burn-in). The bridge acceptance equals the probability that the bridge changes $z$, since an accepted bridge move always flips $z$.

| Analysis | $\widehat{\Pr}(z \text{ changes})$ | UC–AC bridge acceptance |
|---|---|---|
| DENV-4 | 0.423 | 0.503 |
| 31-taxon *rbcL*, JTT | 0.012 | $1.9 \times 10^{-5}$ |
| 31-taxon *rbcL*, OBAMA | 0.100 | $8.1 \times 10^{-3}$ |
| RSV-A | 0.006 | $4.1 \times 10^{-4}$ |
| H3N2 | 0.168 | 0.129 |

#### 4.2 Within-model autocorrelated subtree proposal

The cross-model operators of Section 4.1 address transitions between the UCLN and autocorrelated components, but they do not themselves improve within-component mixing of the branch-rate vector under the autocorrelated component, where the prior induces strong correlations between parent and child rates along the tree. We implemented `ACSubtreeUIncrementOperator` as a block proposal tailored to these correlations. The operator activates only when $z = 1$ and leaves the indicator unchanged, so it is strictly a within-model move.

A subtree root is drawn uniformly from the internal nodes, subject to an upper limit on the number of descendant branches so that the block size stays manageable on large trees. For each branch $b$ in the chosen subtree $S$, the operator reads the current whitened residual under the autocorrelated parameterisation,

$$u_b \;=\; \frac{\log r_b - \log r_{\mathrm{pa}(b)} + \frac{1}{2}\nu_{\mathrm{A}} t_b}{\sqrt{\nu_{\mathrm{A}} t_b}}, \tag{16}$$

perturbs it by an independent Gaussian increment, $u'_b = u_b + \delta\,\epsilon_b$ with $\epsilon_b \sim \mathcal{N}(0,1)$, and reconstructs the new log-rates by pushing the perturbed residuals forward through the autocorrelated map with the same $\nu_{\mathrm{A}}$, the same branch durations, and the boundary parent log-rate held fixed. Ancestors of the subtree root, as well as descendants outside $S$, are left unchanged, so the autocorrelated structure at the subtree boundary is preserved.

Because the state variable is $\boldsymbol{r}$ rather than $\mathbf{u}$, the symmetric Gaussian proposal in $\mathbf{u}$-space induces a change-of-variables correction in log-rate space. The Jacobian of the inverse map $\boldsymbol{r} \mapsto \mathbf{u}$ is triangular when branches are ordered by depth, with diagonal entries $1/(r_b\sqrt{\nu_{\mathrm{A}} t_b})$, and the $\nu_{\mathrm{A}} t_b$ terms cancel between the forward and reverse directions, leaving the log-Hastings correction

$$\log \left| J_{\mathrm{AC}}^{\mathrm{subtree}} \right| \;=\; \sum_{b \in S} \log \frac{r'_b}{r_b}. \tag{17}$$

This construction is a localised variant of the non-centered parameterisation (Papaspiliopoulos et al., 2007) already used by `UCACSwitchBridgeOperator` across the UC/AC boundary, applied instead within the autocorrelated component to allow coordinated updates of correlated rate blocks.

#### 4.3 Within-model non-centered hyperparameter proposals

Two further within-model proposals update the relaxed-clock hyperparameters under the non-centered parameterisation. A vanilla scale proposal on $\sigma_{\mathrm{U}}$ or $\nu_{\mathrm{A}}$ with $\boldsymbol{r}$ held fixed typically proposes a rate configuration that is implausible under the new hyperparameter value, and is therefore almost always rejected on large data sets where the hyperparameter and the branch-rate vector are strongly coupled through the relaxed-clock prior. Both operators co-transform $\boldsymbol{r}$ in lockstep with the hyperparameter so that each branch-rate retains the same quantile position under the relaxed-clock prior.

`UCLDStdevNonCenteredOperator` proposes a move when $z = 0$. It draws $\sigma'_{\mathrm{U}} = \sigma_{\mathrm{U}} \exp(\epsilon)$ with $\epsilon \sim \mathrm{Uniform}(-w, +w)$, holds the UCLN whitened residual $u_b = (\log r_b + \frac{1}{2}\sigma_{\mathrm{U}}^2)/\sigma_{\mathrm{U}}$ fixed for every branch, and rebuilds the log-rates as $\log r'_b = -\frac{1}{2}\sigma_{\mathrm{U}}'^2 + \sigma'_{\mathrm{U}} u_b$. The joint map $(\boldsymbol{r}, \sigma_{\mathrm{U}}) \mapsto (\boldsymbol{r}', \sigma'_{\mathrm{U}})$ has log-Jacobian

$$\log \left| J_{\mathrm{UC}}^{\mathrm{hyp}} \right| \;=\; \sum_{b \in B} \log \frac{r'_b}{r_b} \;+\; (|B| + 1)\,\epsilon, \tag{18}$$

which is the entire log-Hastings correction of the move. The first term is the change-of-variables correction between the rate and log-rate coordinates. In the $(|B|+1)\epsilon$ term, the $|B|$ comes from the branch-rate coordinates: under UCLN $\log r_b = -\frac{1}{2}\sigma_{\mathrm{U}}^2 + \sigma_{\mathrm{U}} u_b$ is linear in $\sigma_{\mathrm{U}}$, so $\partial \log r'_b / \partial \log r_b = \sigma'_{\mathrm{U}}/\sigma_{\mathrm{U}} = e^{\epsilon}$; the $+1$ is the contribution of rescaling the hyperparameter $\sigma_{\mathrm{U}}$ itself.

`ACSigma2NonCenteredOperator` proposes a move when $z = 1$. It draws $\nu'_{\mathrm{A}} = \nu_{\mathrm{A}} \exp(\epsilon)$ with $\epsilon \sim \mathrm{Uniform}(-w, +w)$, holds the autocorrelated whitened residual $u_b$ of Eq. 16 fixed for every branch, and rebuilds the log-rates top-down through the autocorrelated map with the new $\nu'_{\mathrm{A}}$ and a fixed root log-rate. The joint map $(\boldsymbol{r}, \nu_{\mathrm{A}}) \mapsto (\boldsymbol{r}', \nu'_{\mathrm{A}})$ has log-Jacobian

$$\log \left| J_{\mathrm{AC}}^{\mathrm{hyp}} \right| \;=\; \sum_{b \in B} \log \frac{r'_b}{r_b} \;+\; (\tfrac{1}{2}|B| + 1)\,\epsilon, \tag{19}$$

which is likewise the entire log-Hastings correction. As in the UCLN case, the $+1$ is the contribution of the rescaled hyperparameter $\nu_{\mathrm{A}}$ itself; the root log-rate $x_0$ is held fixed and contributes nothing. All $|B|$ non-root branches are updated, but each contributes only $\frac{1}{2}\epsilon$ rather than $\epsilon$, because the autocorrelated map depends on $\nu'_{\mathrm{A}}$ through $\sqrt{\nu'_{\mathrm{A}}}$, giving $\partial \log r'_b / \partial \log r_b = \sqrt{\nu'_{\mathrm{A}}/\nu_{\mathrm{A}}} = e^{\epsilon/2}$. The factor $\frac{1}{2}$ is therefore the exponent of this square-root dependence, not a count of branches.

**Operator tuning and diagnostics.** The within-model non-centered proposals were assigned weights of 15.0 for `ACSubtreeUIncrementOperator`, 15.0 for `UCLDStdevNonCenteredOperator`, and 15.0 for `ACSigma2NonCenteredOperator`. Because the latter three operators are active only in one relaxed-clock component, raw acceptance fractions are not directly comparable across analyses: proposals made while the chain is in the other component are immediate operator rejections. We therefore report conditional acceptance probabilities,

$$\widehat{a}_{\mathrm{cond}} = \frac{N_{\mathrm{accept}}}{N_{\mathrm{accept}} + N_{\mathrm{reject}} - N_{\mathrm{rejectOp}}},$$

where $N_{\mathrm{rejectOp}}$ denotes operator-level rejections, dominated here by proposals attempted while the indicator was in the non-matching state.

The operator weights above were fixed at 15.0 and not tuned per analysis. Each non-centered proposal used a fixed step size, identical across all analyses: a log-normal scale increment with half-width $w = 0.20$ for `UCLDStdevNonCenteredOperator` and $w = 0.15$ for `ACSigma2NonCenteredOperator`, and a residual-increment scale of $\delta = 0.25$ for `ACSubtreeUIncrementOperator`. These step sizes were not adapted during sampling; the conditional acceptance probabilities reported below were used only as a post-hoc mixing diagnostic, not as a tuning target.

Table S4 reports these conditional acceptance probabilities for the three within-model non-centered proposals across all empirical analyses. They lie between roughly 0.5 and 0.98, indicating that, once operator-level rejections from non-matching indicator states are excluded, all three proposals mixed efficiently and none required per-analysis retuning.

Table S4: Conditional acceptance probabilities for the within-model non-centered proposals. For operators active only in one relaxed-clock component, conditional acceptance was computed after excluding operator-level rejections caused by attempts in the non-matching indicator state. Numbers in parentheses give the resulting number of valid proposals.

| Analysis | `ACSubtreeUIncrementOperator` | `UCLDStdevNonCenteredOperator` | `ACSigma2NonCenteredOperator` |
|---|---|---|---|
| DENV-4 | 0.716 ($n$ = 299,887) | 0.876 ($n$ = 527,707) | 0.933 ($n$ = 299,815) |
| RSV-A | 0.761 ($n$ = 188) | 0.548 ($n$ = 874,313) | 0.983 ($n$ = 175) |
| 31-taxon *rbcL*, JTT | 0.519 ($n$ = 691,593) | 0.610 ($n$ = 15,413) | 0.830 ($n$ = 691,861) |
| 31-taxon *rbcL*, OBAMA | 0.517 ($n$ = 43,369) | 0.640 ($n$ = 54,776) | 0.678 ($n$ = 43,944) |
| H3N2 | 0.716 ($n$ = 15,862,192) | 0.860 ($n$ = 8,406,254) | 0.782 ($n$ = 6,800,792) |

### 5 Supplementary Note 5. Branch-level diagnostics across empirical analyses

We generated branch-level diagnostic plots for empirical analyses whose posterior tree logs contained branch-rate metadata. These diagnostics are not all interpreted in the same way. When posterior support was concentrated on the strict clock or on a single relaxed-clock family, UCLN-versus-autocorrelated branch-rate comparisons were treated as diagnostics rather than as primary branch-level summaries. We interpret UCLN-versus-autocorrelated branch-rate comparisons only when both relaxed-clock families retained non-negligible posterior support.

Table S5 summarises the status and interpretation of these diagnostics across the empirical analyses. The H3N2 posterior tree logs contained structured-coalescent type annotations but did not contain branch-rate metadata, so analogous branch-rate diagnostics could not be reconstructed from the archived H3N2 tree samples without rerunning or resuming the analysis with branch-rate logging enabled.

Table S5: Status and interpretation of branch-level diagnostics across empirical analyses.

| Analysis | Clock-support pattern | Diagnostic status | Interpretation |
|---|---|---|---|
| DENV-4 | Strict clock received strongest posterior support | CCD0-MAP tree shown; UC–AC scatter generated and archived | CCD0 tree is treated as a branch-level diagnostic. UC–AC scatter is not interpreted as a primary branch-rate summary because the main posterior support was not concentrated within the relaxed class. |
| *rbcL*, JTT | Autocorrelated dominated | CCD0-MAP tree shown; UC–AC scatter generated and archived | CCD0 tree shows an effectively autocorrelated-clock-dominated rate-multiplier pattern. UC–AC scatter is not interpreted because UCLN support was minor ($P_{\mathrm{UC}} = 0.032$). |
| *rbcL*, OBAMA | UCLN and autocorrelated both supported | UC–AC scatter shown in the main text; CCD0-MAP tree shown here | Relaxed-clock branch-rate comparison is interpretable because both relaxed-clock families retained substantial posterior support. |
| RSV-A | UCLN dominated | CCD0-MAP tree shown in the main text; UC–AC scatter generated and archived | CCD0 tree is interpreted as UCLN-dominated. UC–AC scatter is not interpreted because $P_{\mathrm{AC}} < 0.01$. |
| H3N2 | Relaxed class supported; autocorrelated dominant | Not shown | Archived tree logs contained type annotations but no branch-rate metadata, so branch-rate diagnostics would require a resumed or repeated run with branch-rate logging enabled. |

#### 5.1 CCD0-MAP rate-coloured trees

For DENV-4, branch colours are on the absolute substitution-rate scale, in substitutions site$^{-1}$ year$^{-1}$. For the *rbcL* analyses, branch colours represent relative branch-rate multipliers, which are unitless. The baseline JTT and OBAMA *rbcL* trees were summarised separately from their corresponding posterior tree samples and should not be interpreted as the same fixed topology re-coloured under two site models. Because topology was estimated in both analyses, changing the amino-acid site model can change the posterior tree distribution and hence the CCD0-MAP summary topology.

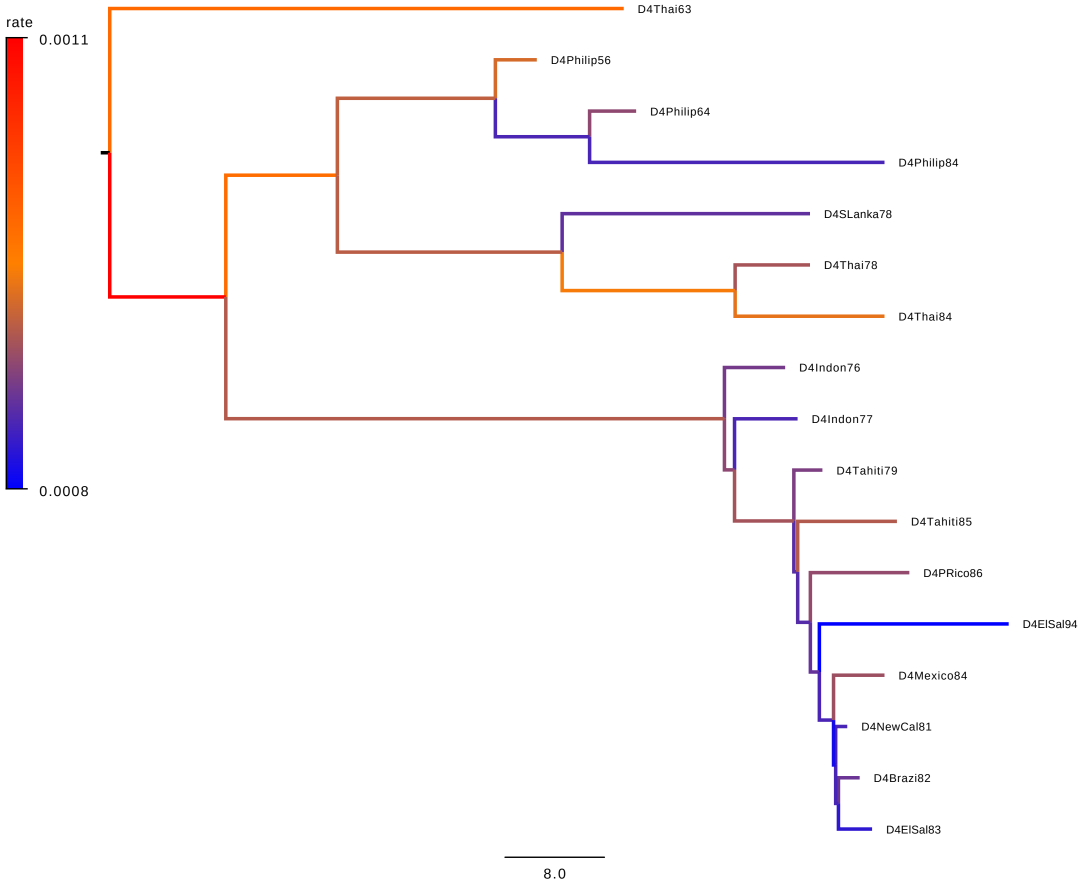

Figure S2: CCD0-MAP summary tree for the DENV-4 benchmark, with branches coloured by posterior branch-rate summaries from the hierarchical clock-mixture analysis. Colour-bar values are in substitutions site$^{-1}$ year$^{-1}$, and branch lengths are in years. Because the DENV-4 benchmark placed strongest posterior support on the strict clock, this rate-coloured tree is included as a branch-level diagnostic rather than as a primary model-averaged branch-rate summary.

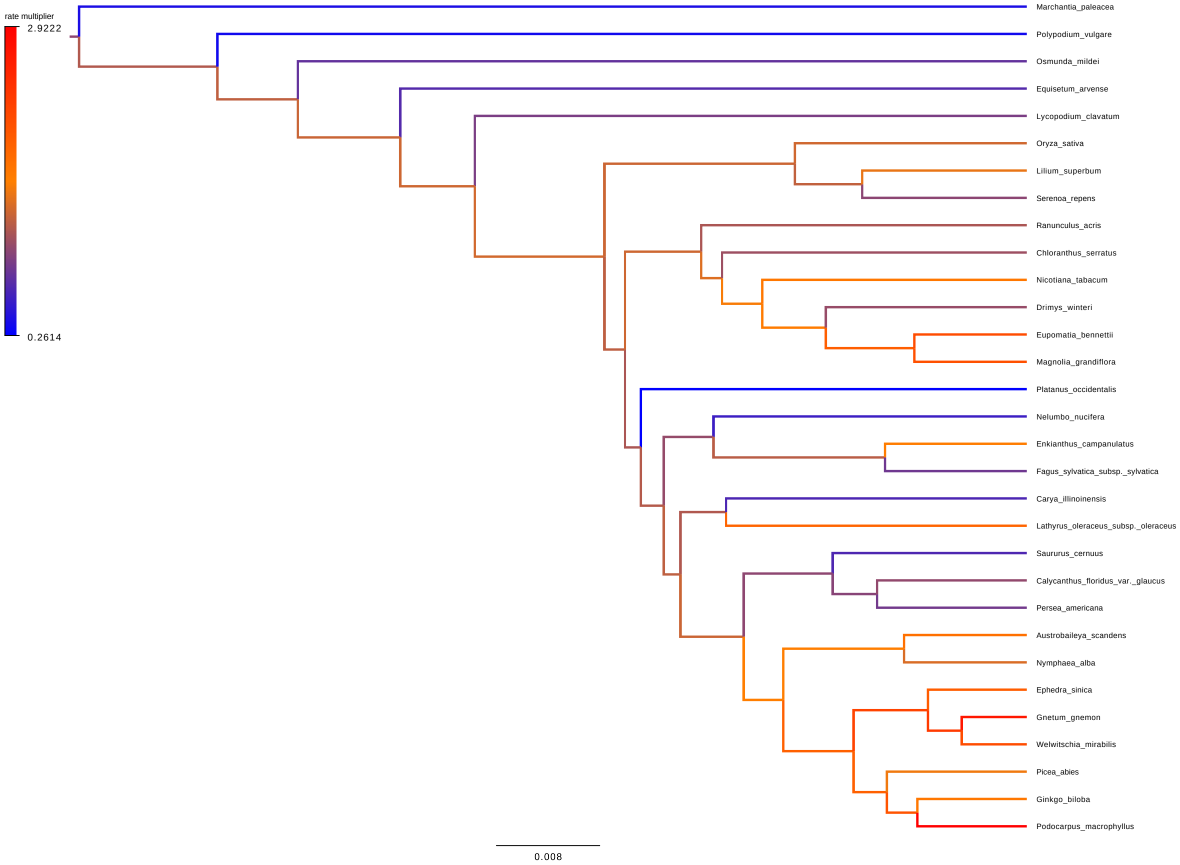

Figure S3: CCD0-MAP summary tree for the baseline *rbcL* JTT analysis. Branches are coloured by posterior relative branch-rate multipliers from the hierarchical clock-mixture analysis. Colour-bar values are unitless. Because posterior support under the JTT analysis was concentrated almost entirely on the autocorrelated clock, the displayed branch-rate pattern is effectively autocorrelated-clock dominated.

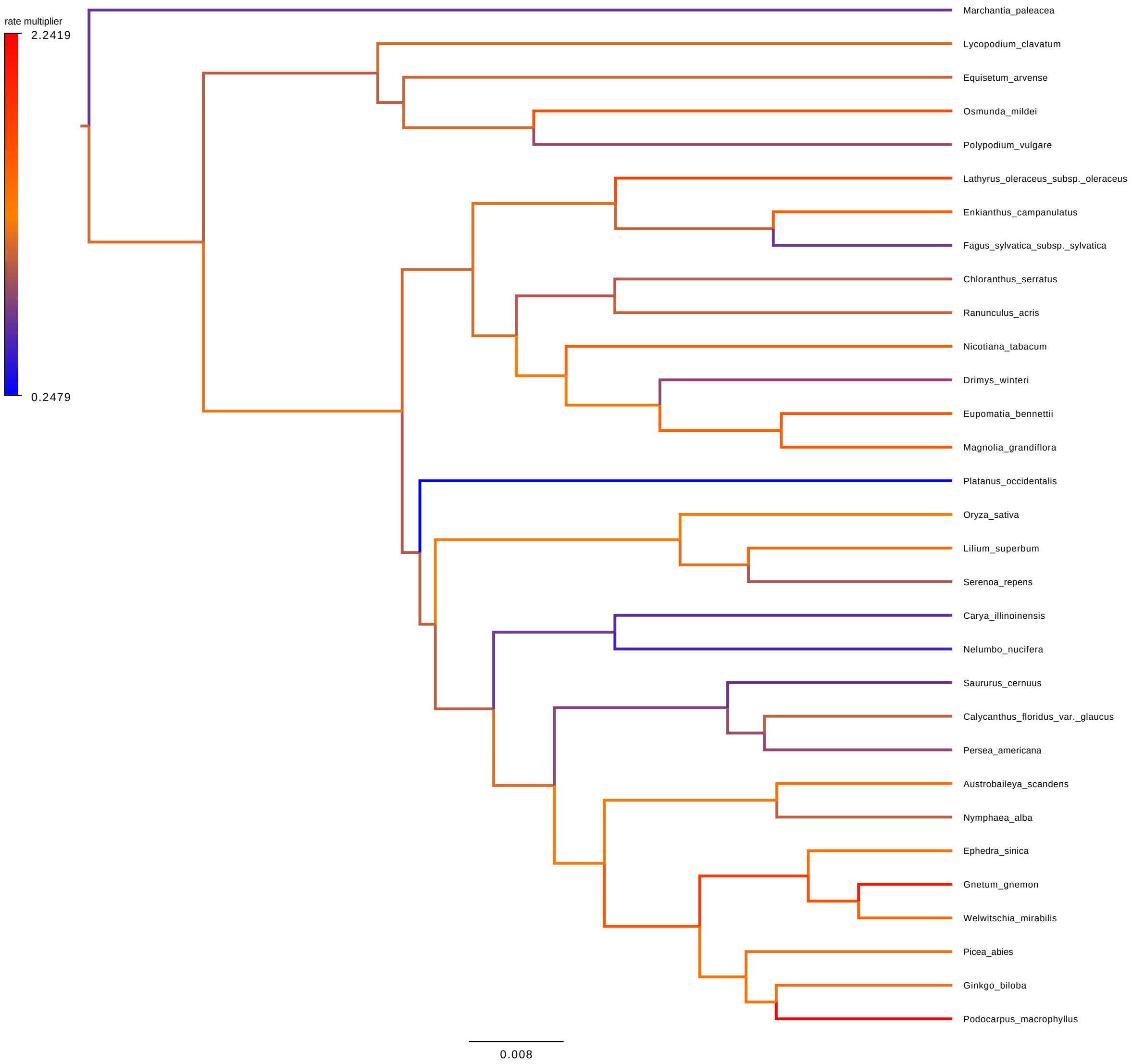

Figure S4: CCD0-MAP summary tree for the *rbcL* OBAMA site-model sensitivity analysis. Branches are coloured by posterior relative branch-rate multipliers from the hierarchical clock-mixture analysis. Colour-bar values are unitless. This tree was summarised separately from the OBAMA posterior tree sample and is therefore not constrained to have the same topology as the baseline JTT CCD0-MAP tree.
